# Trans-omic analysis reveals allosteric and gene regulation-axes for altered glucose-responsive liver metabolism associated with obesity

**DOI:** 10.1101/653758

**Authors:** Toshiya Kokaji, Atsushi Hatano, Yuki Ito, Katsuyuki Yugi, Miki Eto, Satoshi Ohno, Masashi Fujii, Ken-ichi Hironaka, Riku Egami, Hiroshi Inoue, Shinsuke Uda, Hiroyuki Kubota, Yutaka Suzuki, Kazutaka Ikeda, Makoto Arita, Masaki Matsumoto, Keiichi I. Nakayama, Akiyoshi Hirayama, Tomoyoshi Soga, Shinya Kuroda

**Affiliations:** Department of Biological Sciences, Graduate School of Science, University of Tokyo, 7-3-1 Hongo, Bunkyo-ku, Tokyo 113-0033, Japan; YCI Laboratory for Trans-Omics, Young Chief Investigator Program, RIKEN Center for Integrative Medical Science, 1-7-22 Suehiro-cho, Tsurumi-ku, Yokohama, Kanagawa 230-0045, Japan; Department of Computational Biology and Medical Sciences, Graduate School of Frontier Sciences, University of Tokyo, 5-1-5 Kashiwanoha, Kashiwa, Chiba 277-8562, Japan; Institute for Advanced Biosciences, Keio University, Fujisawa, 252-8520, Japan; PRESTO, Japan Science and Technology Agency, 1-7-22 Suehiro-cho, Tsurumi-ku, Yokohama, Kanagawa 230-0045, Japan; Molecular Genetics Research Laboratory, Graduate School of Science, University of Tokyo, 7-3-1 Hongo, Bunkyo-ku, Tokyo 113-0033, Japan; Program of Mathematical and Life Sciences, Graduate School of Integrated Sciences for Life, Hiroshima University, Kagamiyama, Higashi-hiroshima 739-8526, Japan; Metabolism and Nutrition Research Unit, Institute for Frontier Science Initiative, Kanazawa University, 13-1 Takaramachi, Kanazawa, Ishikawa, 920-8641, Japan; Division of Integrated Omics, Research Center for Transomics Medicine, Medical Institute of Bioregulation, Kyushu University, 3-1-1 Maidashi, Higashi-ku, Fukuoka 812-8582, Japan; Laboratory for Metabolomics, RIKEN Center for Integrative Medical Sciences, Yokohama, Japan; Graduate School of Medical Life Science, Yokohama City University, Yokohama, Japan; Division of Physiological Chemistry and Metabolism, Keio University Faculty of Pharmacy, Tokyo, Japan; Department of Molecular and Cellular Biology, Medical Institute of Bioregulation, Kyushu University, 3-1-1 Maidashi, Higashi-ku, Fukuoka 812-8582, Japan; Institute for Advanced Biosciences, Keio University, 246-2 Mizukami, Kakuganji, Tsuruoka, Yamagata 997-0052, Japan; Core Research for Evolutional Science and Technology (CREST), Japan Science and Technology Agency, Bunkyo-ku, Tokyo 113-0033, Japan

## Abstract

Impaired glucose tolerance associated with obesity causes postprandial hyperglycemia and can lead to type 2 diabetes. To study the differences in liver metabolism in the healthy and obese states, we constructed and analyzed trans-omic glucose-responsive metabolic networks with layers for metabolites, expression data for metabolic enzyme genes, transcription factors, and insulin signaling proteins from the livers of healthy and obese mice. We integrated multi-omic time-course data from wild-type (WT) and leptin-deficient obese (*ob*/*ob*) mice after orally administered glucose. In WT mice, metabolic reactions were rapidly regulated (within 10 minutes of oral glucose administration) primarily by glucose-responsive metabolites, especially ATP and NADP+, which functioned as allosteric regulators and substrates of metabolic enzymes, and by Akt-dependent glucose-responsive genes encoding metabolic enzymes. In *ob*/*ob* mice, most rapid regulation by glucose-responsive metabolites was absent; instead, glucose administration produced slow changes in the expression of metabolic enzyme-encoding genes to alter metabolic reactions in a time scale of hours. Few common regulatory events occurred in both the healthy and obese mice. Thus, our trans-omic network analysis revealed regulation of liver metabolism in response to glucose is mediated through different mechanisms in the healthy and obese states: Rapid changes in allosteric regulators and substrates and in gene expression dominate the healthy state, and slow transcriptional regulation dominates the obese state.

**One Sentence Summary:** Rapid changes in regulatory metabolites and gene expression dominate the healthy state, and slow transcriptional regulation dominates the obese state.

---

The ability to produce stable blood glucose is indispensable for human life and health (*1*–*3*). Although a large amount of glucose enters the body through meals, changes in organ metabolism maintains glucose homeostasis (*4*–*6*). Impairment of the regulation of organ metabolism, commonly due to obesity and insulin resistance, results in hyperglycemia and development of type 2 diabetes mellitus (*4*–*6*). The liver, into which dietary glucose flows directly through the portal vein, has a primary function in maintaining glucose homeostasis (*7, 8*). Indeed, the liver is both a glucose-producing organ, supplying glucose for extra-hepatic organs, and glucose-utilizing organ, metabolizing one third of orally administered glucose (*8, 9*). Oral intake of glucose produces drastic changes in the liver metabolism— not only glucose metabolism but also lipid and amino-acid metabolism, collectively glucose-responsive metabolism. The mechanisms regulating glucose-responsive metabolism in the liver and how these mechanisms are altered in obesity have yet to be identified.

Metabolism is a set of chemical reactions that convert one metabolite into another. Chemical reactions in metabolism, denoted here as metabolic reactions, involve metabolites, which function as substrates, products, and allosteric regulators of the metabolic enzymes that catalyze the reactions. Metabolic reactions are also regulated by cellular processes that affect the amount of the enzyme (for example, changes in gene expression) or posttranslational modifications of the enzymes (for example, phosphorylation). The amount and phosphorylation status of metabolic enzymes are regulated by transcription factors that control gene expression and signaling molecules that control transcription factor activity and metabolic enzyme activity through phosphorylation. Therefore, metabolic reactions are regulated by an integrated network consisting of metabolites, the metabolic enzymes and their phosphorylation status, transcription factors and their activation status, and signaling molecules that mediate phosphorylation of the metabolic enzymes and the transcription factors. We propose that regulatory mechanisms controlling metabolic reactions in the liver and the alterations in these processes that are associated with obesity can be investigated by integrating simultaneous measurements of metabolite abundance, expression of genes for and phosphorylation status of metabolic enzymes and transcription factors, and the abundance and activation status of signaling molecules into a multi-layered trans-omic network.

Omic measurements, such as metabolomic, transcriptomic, and proteomic measurements, enable large-scale measurement of molecules in each layer of the multi-omic network (*10*–*14*). We have applied an approach that we call ‘trans-omic’ analysis for the construction of global biochemical network using simultaneously measured multi-omic data based on direct molecular interaction (*15*–*17*). We used this approach to construct an insulin-induced regulatory trans-omic network for metabolism of FAO hepatoma cells by integrating simultaneously measured multi-omic measurements (*15, 18*). With this trans-omic network, we discovered selective regulation of the trans-omic network that depended on doses of insulin. However, the regulatory trans-omic network for glucose-responsive metabolic reactions of the liver and the alterations in this regulatory network that are associated with obesity have yet to be identified.

Here, to reveal the global regulatory mechanisms controlling glucose-responsive metabolism in the liver in the healthy and obese states, we constructed regulatory trans-omic networks for glucose-responsive metabolic reactions in the liver of wild type (WT) and leptin-deficient obese (*ob*/*ob*) mice. We administered glucose to the mice orally and then obtained simultaneously measured multi-omic data, which we integrated into a trans-omic network. The *ob*/*ob* mice are a widely used model of obesity and insulin resistance, because these mice become profoundly obese by overeating due to deficiency of the anorexigenic hormone leptin (*19*). We found that, in WT mice, glucose-responsive metabolic reactions were mainly regulated by rapid changes in metabolites, which function as allosteric regulators and substrates of metabolic enzymes, and by rapid changes in Akt-dependent gene expression of metabolic enzymes. By contrast, in *ob*/*ob* mice, most of the rapid changes in metabolites were absent. Instead, metabolic reactions were regulated by a slower change in gene expression, which was not seen in WT mice. Thus, we identified profoundly different mechanisms of glucose-responsive regulation of metabolic reactions in the liver in the healthy and obese states.

## Results

### The pipeline of the construction of regulatory trans-omic network for glucose-responsive metabolic reactions

In this study, we constructed the regulatory trans-omic networks for glucose-responsive metabolic reactions in the liver of WT and *ob*/*ob* mice, and examined differences in regulatory mechanisms controlling these reactions (Fig. 1). Metabolic reactions are regulated by an integrated network consisting of (i) metabolites as allosteric regulators, substrates, and products, (ii) metabolic enzymes, (iii) transcription factors, and (iv) signaling molecules (fig. S1). Here, we measured metabolite abundance, gene expression, and protein abundance and phosphorylation of signaling molecules by multi-omic analyses, enzymatic assays, and Western blotting in the liver and blood of WT and *ob*/*ob* mice following oral glucose administration. We used these data and bioinformatic resources to construct regulatory trans-omic networks for glucose-responsive metabolic reactions of the liver in WT and *ob*/*ob* mice (Fig. 1).

**Figure 1.**
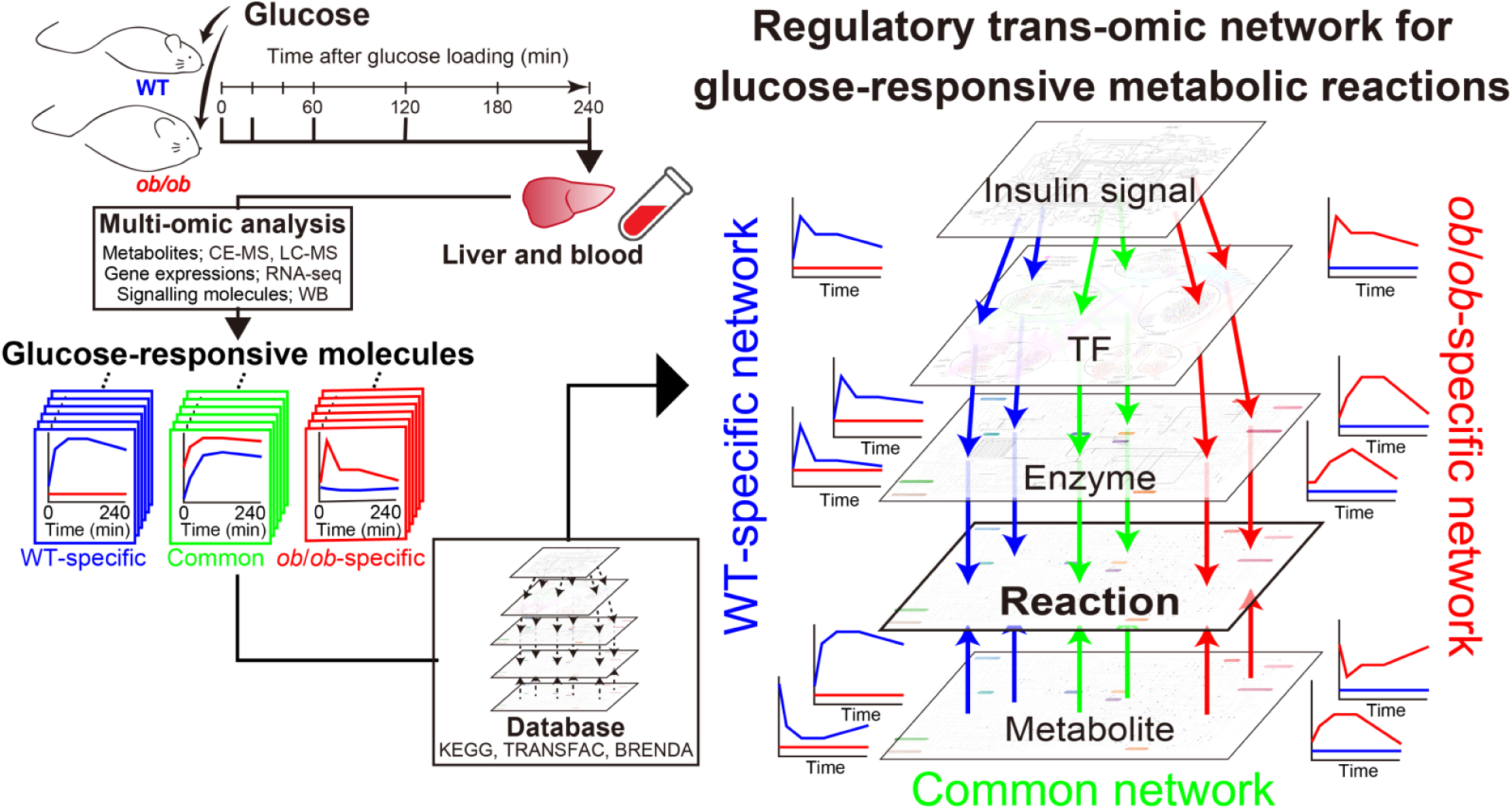
The pipeline of the construction of regulatory trans-omic network for glucose-responsive metabolic reactions. We measured the time courses of multi-omic data from the liver and blood of WT mice and *ob*/*ob* mice following oral glucose administration and identified molecules that changed by oral glucose administration, which we defined as glucose-responsive molecules in each layer (see Materials and Methods: Identification of glucose-responsive molecules). We added inter-layer regulatory connections between glucose-responsive molecules in different layers using bioinformatic methods and information in public databases. The result was a regulatory trans-omic network for glucose-responsive metabolic reactions in the liver of WT and *ob*/*ob* mice. We identified regulatory trans-omic subnetworks specific to WT mice (blue), *ob*/*ob* mice (red), and common to both mice (green).

We orally administered glucose to 16 h-fasting WT and *ob*/*ob* mice, sacrificed the animals, and collected the liver and blood at 0, 20, 60, 120, 240 min after administration (fig. S2A). We analyzed the liver samples by performing metabolomics, transcriptomics, and Western blotting, and analyzed blood samples by performing metabolomics (fig. S2B). The liver samples provided data for phosphorylation-mediated changes in signaling molecules, transcription factors, and enzymes involved in insulin signaling (Western blotting data), the gene expression information of transcription factors and enzymes (transcriptomic data), and metabolite abundance information (metabolomic data). We defined “glucose-responsive molecules” as the molecules that were quantitatively changed by oral glucose administration. Metabolites and genes that showed an absolute log_2_ fold change larger than 0.585 (2^0.585^ = 1.5) and an FDR-adjusted p value (q value) less than 0.1 at any time point compared to the fasting state (0 min) were defined as glucose-responsive. To investigate differences in the regulatory networks, we focused on the relative glucose-responsiveness of molecules, meaning the change from fasting state, in the WT and *ob*/*ob* mice, rather than the differences in the absolute amounts of molecules, which are described in Supplementary Text: The differences of the amounts of molecules between WT mice and *ob*/*ob* mice before oral glucose administration.

We integrated the glucose-responsive molecules into a multi-omic layered network, and identified the regulatory connections among the layers to generate the regulatory trans-omic networks for glucose-responsive metabolic reactions of the liver in WT and *ob*/*ob* mice. The regulatory trans-omic network consisted of layers of insulin signaling molecules (Insulin signal), transcription factors (TF), the gene expression and phosphorylation of metabolic enzyme (Enzyme), metabolic reactions (Reaction), metabolites (Metabolite), and the regulatory connections between the layers (Fig. 1). By comparing the regulatory trans-omic networks, we identified common elements in the glucose-responsive liver metabolic networks between WT and *ob/ob* mice, which we referred to as the “common network” and show with green, as well as identified glucose-responsive regulatory networks distinct to the healthy (WT-specific network, blue) or obese states (*ob/ob*-specific network, red).

### Identification of glucose-responsive metabolites

We measured metabolomic changes in the liver of WT and *ob*/*ob* mice following oral glucose administration using capillary electrophoresis mass spectrometry (CE-MS) and liquid chromatography mass spectrometry (LC-MS) (Fig. 2, figs. S3 and S4, tables S1 and S2, and Supplementary Text). Using CE-MS, we quantified 161 polar metabolites, including carbohydrates, amino acids, and nucleic acids; using LC-MS, we quantified 15 lipids; and, using enzymatic assays, we quantified glycogen, a polar metabolite, and triglyceride, a lipid. We identified glucose-responsive metabolites in the liver of WT and *ob*/*ob* mice that exhibited statistically significant changes in response to oral glucose administration (Fig. 2). Many metabolites were not glucose responsive: Their abundance did not change statistically significantly from the fasting values (table S1). We categorized statistically significant changes into increased and decreased groups. To define an increase or decrease in time courses with changes in both directions at different time points, we used the direction of change compared to time 0 at the earliest time point that showed a significant change. We also analyzed metabolites in mice given an equivalent amount of water orally, which showed no changes except for isethionate in *ob/ob* mice (table S1), confirming that the changes we detected reflected a physiological response to the orally administered glucose.

**Figure 2.**
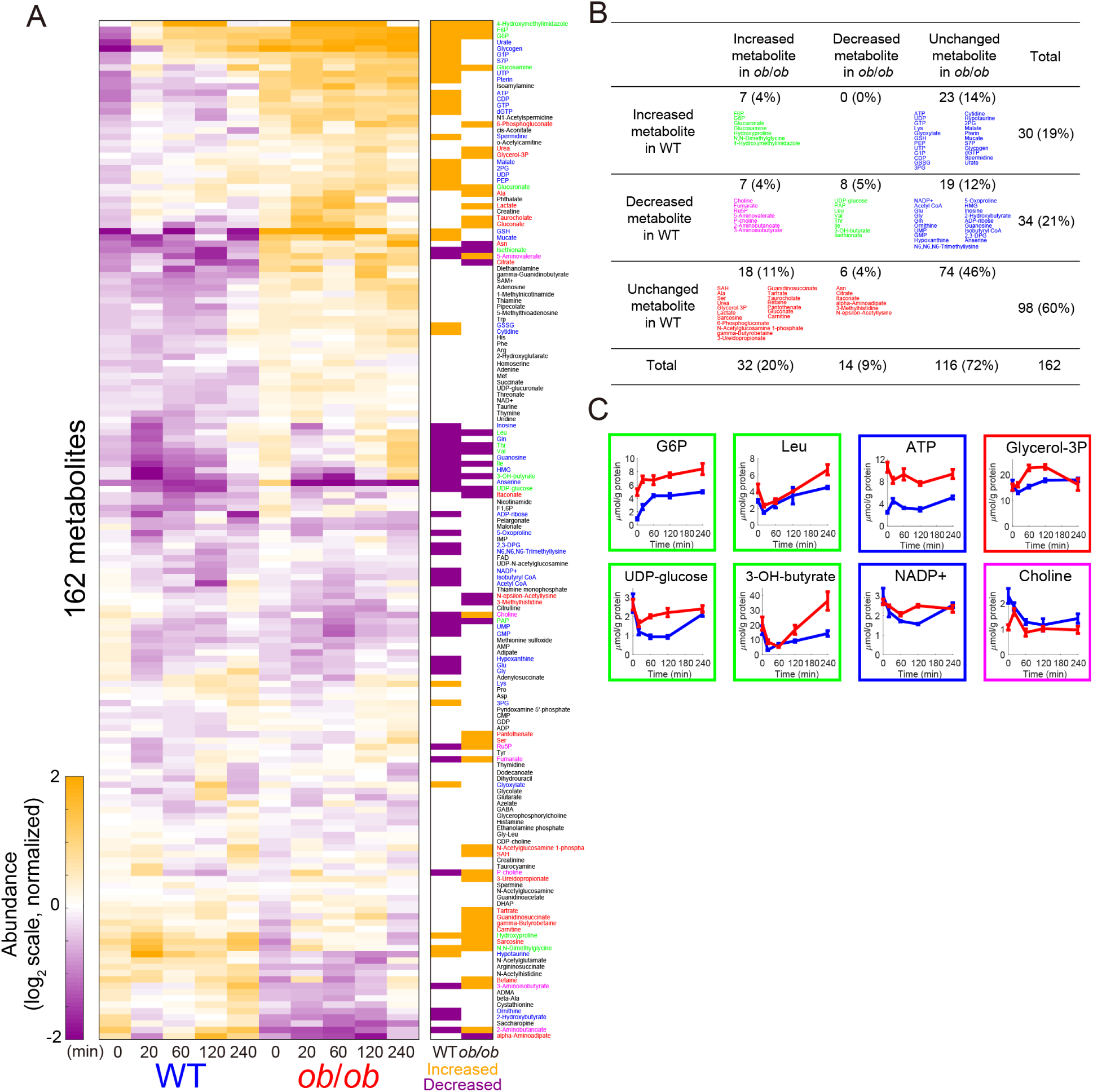
Identification of glucose-responsive metabolites. (**A**) Left: The heat map of the time courses of 162 metabolites from the liver of WT and *ob*/*ob* mice following oral glucose administration. To investigate the changes from fasting state, two time courses of each metabolite were divided by the geometric mean of the values of WT mice and *ob*/*ob* mice in fasting state (0 min), and then log_2_-transformed. Metabolites were ordered by hierarchical clustering using Euclidean distance and Ward’s method (see fig. S3 for characteristics of each cluster and table S1 for data). Right: The bars in the heat map are colored according to the glucose-responsiveness, meaning the change from fasting state (0 min), in the WT and *ob/ob* mice. Metabolites that showed an absolute log_2_ fold change larger than 0.585 (2^0.585^ = 1.5) and an FDR-adjusted p value (q value) less than 0.1 at any time point were defined as glucose-responsive: increased (orange), decreased (purple), or were unchanged (white). To define an increase or decrease in time courses with changes in both directions at different times, we used the direction of change compared to time 0 at the earliest time point that showed a significant change. Metabolites written in red text indicate glucose-responsive metabolites specific to WT; blue text, specific to *ob*/*ob* mice; green text, common to both; pink text, opposite responses between WT mice and *ob*/*ob* mice; black text, no response to glucose. See table S1 for unabbreviated names of metabolites. (**B**) Increased, decreased, and unchanged metabolites in the liver of WT mice and *ob*/*ob* mice: blue text, WT specific; red text, *ob*/*ob* specific; green text, glucose-responsive metabolites common to both; pink text, opposite responses between WT mice and *ob*/*ob* mice. The number of each type of glucose-responsive metabolites and their percentages out of the total quantified metabolites are shown. (**C**) Graphs showing the metabolites with responses that were common to both WT and ob/ob (green boxes), specific to WT mice (blue boxes), and specific to *ob/ob* mice (red boxes), and the metabolites that change in opposite directions in WT and *ob/ob* mice (pink boxes). Within the graphs, blue lines are the responses of the WT mice and red lines are the responses of the *ob/ob* mice. Data are shown as the mean and SEM of 5 mice.

Of the 162 polar metabolites, we determined that a small proportion of them increased or decreased in both WT and *ob/ob* mice: 7 increased in both WT and *ob*/*ob* mice and 8 other metabolites decreased (Fig. 2B, table S1). Glucose-6-phosphate (G6P), the product of first step of glycolysis, increased in both WT and *ob/ob* mice (Fig. 2C). Even though the absolute amounts of G6P differed between WT and *ob*/*ob* mice, G6P was a glucose-responsive metabolite that increased in both. Metabolites that decreased in both mice included the amino acids isoleucine (Ile), leucine (Leu), and valine (Val), the nucleotide sugar UDP-glucose, and the ketone body 3-hydroxybutyrate (3-OH-butyrate) (Fig. 2C). Although Leu had the significantly increased and decreased time points in both, it was categorized as a decreased metabolite because it decreased at the earliest time point. Metabolites that increased only in WT mice included ATP; metabolites that decreased only in WT mice included NADP+ (Fig. 2C). We also measured the time courses of lipidomic changes in the liver of WT and *ob*/*ob* mice, but no lipids showed significant changes in response to oral glucose administration (fig. S4, table S2).

Our metabolomic analysis revealed that the number of glucose-responsive metabolites specific to WT mice (42 = 23 increased + 19 decreased) was larger than that specific to *ob*/*ob* mice (24 = 18 increased + 6 decreased) and that only a limited number were common to both mice (15 = 7 increased + 8 decreased). ATP and NADP+ are notable WT-specific glucose-responsive metabolites because these two cofactors are involved in hundreds of metabolic reactions. This result suggested that ATP and NADP+ regulate many glucose-responsive metabolic reactions in WT mice and that the lack of glucose responsiveness in these two metabolites in *ob/ob* mice could produce a large difference in the regulatory mechanisms between the healthy and obese states.

We also measured the metabolomic changes in the blood of WT and *ob*/*ob* mice, and identified glucose-responsive metabolites in the blood (fig. S5, table S3, and Supplementary Text). Most metabolites in the blood did not show a significant change in response to oral glucose administration. Only nicotinamide, proline, and threonine changed specifically in the blood of WT mice; all increased. Only alpha-aminoadipate, 2’-deoxycytidine, and glycerol-3-phosphate changed specifically in the blood of *ob/ob* mice (fig. S5A), and none showed a change in common to both. In addition, only the changes in 3-OH-butyrate, Val, Leu, Ile, 3-Hydroxy-3-methylglutarate (HMG), and creatine were highly correlated between the blood and the liver in both WT and *ob*/*ob* mice and the other metabolites did not show high correlations (fig. S5B), indicating that contamination of blood in the liver samples had a negligible effect on the liver data.

### Identification of glucose-responsive genes and inference of regulatory connections between TFs and genes

We used RNA sequencing (RNA-seq) to measure the time courses of transcriptomic changes in 14,292 genes in the liver of WT and *ob*/*ob* mice following oral glucose administration (Fig. 3, fig. S6, Table 1, and tables S4 and S5). We identified glucose-responsive genes in the liver of WT and *ob*/*ob* mice (Fig. 3) and performed pathway enrichment analysis of these genes (Table 1 and table S5). Our transcriptomic analysis revealed that the number of *ob*/*ob* mice-specific glucose-responsive genes is larger than that of WT mice-specific glucose-responsive genes: *ob*/*ob*, 1933 = 791 (increased) + 1142 (decreased); WT, 677 = 327 (increased) + 350 (decreased). To confirm that the glucose-responsive genes show smaller changes or no changes to water administration, we measured the expression of a subset of glucose-responsive genes encoding enzymes involved in glycolysis, gluconeogenesis, lipid synthesis, and cholesterol synthesis (table S6). Except for a decrease of genes involved in cholesterol metabolism in *ob*/*ob* mice (fig. S7, A and B), we did not observe significant changes in response to water. Of the glucose-responsive genes, we placed glucose-responsive genes encoding metabolic enzymes in the Enzyme layer of the trans-omic network, and those encoding transcription factors in the TF layer (see Fig. 5).

**Table 1.** Pathway enrichment analysis of the glucose-responsive genes. Pathways with p value < 0.001 are shown.

|  | Upregulated gene in *ob/ob* |  | Downregulated gene in *ob/ob* |  | Unchanged gene in *ob/ob* |  |
| --- | --- | --- | --- | --- | --- | --- |
|  | activity | p value | activity | p value | activity | p value |
| Upregulated gene in WT | p53 signaling pathway | $3.2 \times 10^{-5}$ | Terpenoid backbone biosynthesis | $2.8 \times 10^{-9}$ | Metabolic pathways | $9.0 \times 10^{-9}$ |
| | | | Steroid biosynthesis | $1.8 \times 10^{-8}$ | Steroid biosynthesis | $2.2 \times 10^{-5}$ |
| | | | Metabolic pathways | $1.9 \times 10^{-4}$ | Drug metabolism - cytochrome P450 | $1.7 \times 10^{-4}$ |
| | | | | | Glutathione metabolism | $5.3 \times 10^{-4}$ |
| | | | | | Retinol metabolism | $9.0 \times 10^{-4}$ |
| Downregulated gene in WT | Retinol metabolism | $1.3 \times 10^{-4}$ | | | Linoleic acid metabolism | $2.9 \times 10^{-6}$ |
| | Steroid hormone biosynthesis | $8.9 \times 10^{-4}$ | | | Arachidonic acid metabolism | $3.5 \times 10^{-6}$ |
| | Linoleic acid metabolism | $9.6 \times 10^{-4}$ | | | Retinol metabolism | $2.5 \times 10^{-4}$ |
| Unchanged gene in WT | | | Ribosome | $1.6 \times 10^{-16}$ | | |
| | | | Phagosome | $2.6 \times 10^{-4}$ | | |

**Figure 3.**
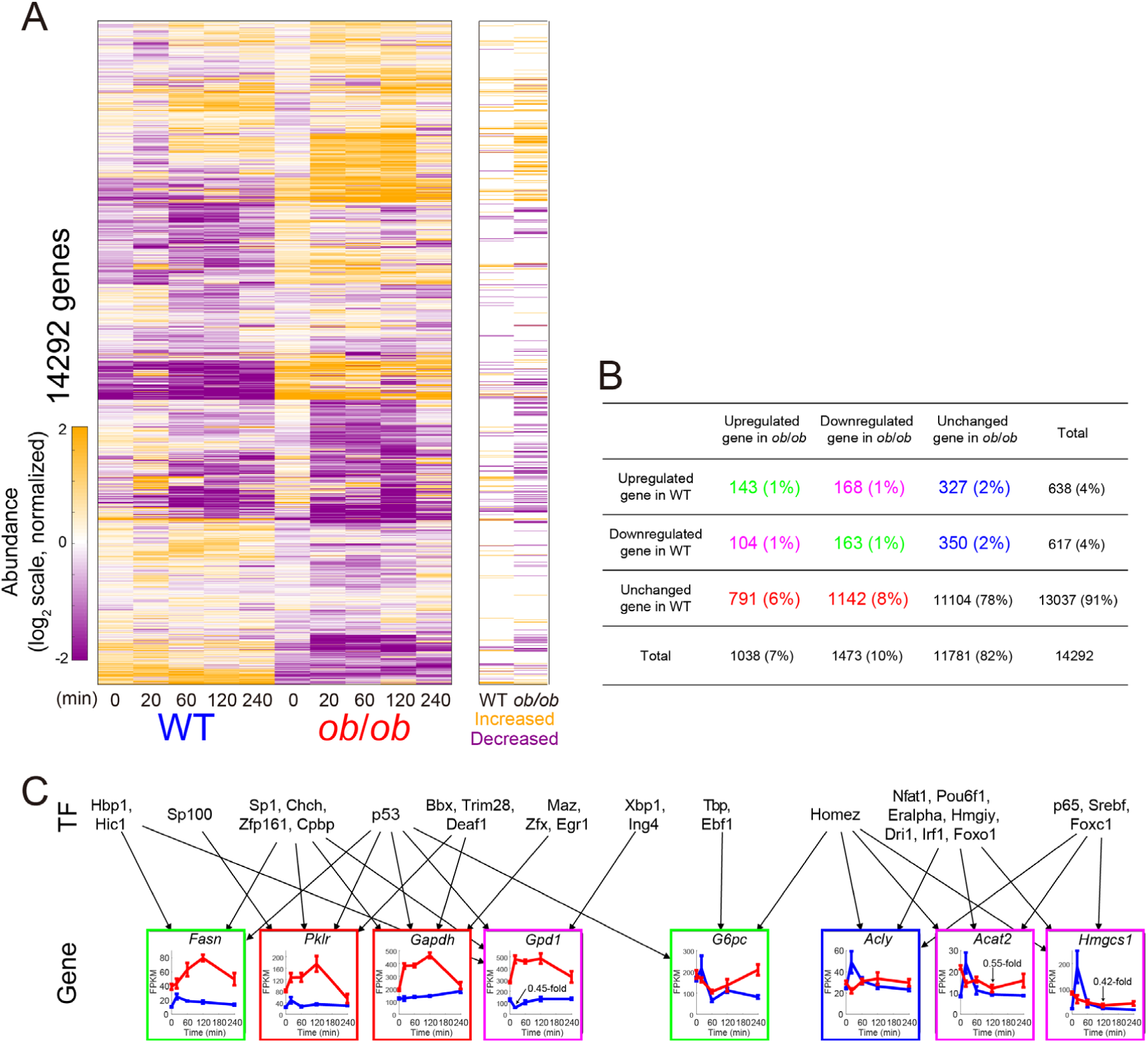
Identification of glucose-responsive genes. (**A**) Left: The heat map of the time courses of transcript abundance for 14,292 genes in the livers of WT and *ob*/*ob* mice following oral glucose administration. To investigate the changes from fasting state, two time courses of each metabolite were divided by the geometric mean of the values of WT mice and *ob*/*ob* mice in fasting state (0 min), and then log_2_-transformed. Genes were ordered by hierarchical clustering using Euclidean distance and Ward’s method (see table S4 for data). Right: The bars in the heat map are colored according to the glucose responsiveness, meaning the change from fasting state (0 min), in the WT and *ob/ob* mice. Genes that showed an absolute log_2_ fold change larger than 0.585 (2^0.585^ = 1.5) and a q value less than 0.1 at any time point were defined as glucose-responsive: increased (orange), decreased (purple), or were unchanged (white). (**B**) Upregulated, downregulated, and unchanged genes in the liver of WT mice (row) and *ob*/*ob* mice: blue, WT specific; red, *ob*/*ob* specific; green, glucose-responsive metabolites common to both; pink, opposite responses between WT mice and *ob*/*ob* mice. The number of each type of glucose-responsive genes and their percentages out of the total quantified genes are shown. (**C**) Graphs showing the gene expression time courses for the indicated genes. Genes include those that exhibited changes in common to both WT and *ob/ob* (green boxes), changes specific to WT mice (blue boxes), changes specific to *ob/ob* mice (red boxes), and changes in opposite directions in each (pink boxes). Within the graphs, blue lines are the responses of the WT mice and red lines are the responses of the *ob/ob* mice. Data are shown as the mean and SEM of mice (n = 11 or 12 at 0 min, n = 3 at 20 min, n = 3 at 60 min, n = 3 at 120 min, and n = 3 at 240 min). The inferred regulatory connections are shown as arrows from transcription factors to genes. The regulatory connections were inferred using hierarchical clustering analysis of gene expression time courses together with a transcription factor database TRANSFAC (*21, 22*). See fig. S6 for statistical confidence in inferred transcription factors; see table S7 for the unabbreviated names of the transcription factors.

Similar to the metabolites, we found that only a small proportion of genes were regulated in common between the two states: 143 genes (1% of the total quantified genes) were upregulated and 163 (1%) were downregulated in both (Fig. 3B). Genes upregulated in common included fatty acid synthase (*Fasn*) (Fig. 3C). Downregulated genes common to both mice included the gene encoding glucose-6-phosphatase (*G6pc*), a key metabolic enzyme of gluconeogenesis (*20*) (Fig. 3C).

Pathway enrichment analysis showed that the 327 genes that were specifically upregulated in WT mice were enriched for enzymes involved in steroid biosynthesis (Table 1), indicating an increase in cholesterol biosynthesis. Furthermore, 168 genes that were upregulated in WT mice and downregulated in *ob*/*ob* mice were enriched for enzymes involved in steroid biosynthesis and terpenoid backbone synthesis, such as 3-hydroxy-3-methylglutaryl CoA synthase 1 (*Hmgcs1*) and acyl-CoA: cholesterol acyltransferase 2 (*Acat2*) (Fig. 3C, pink boxes). The responses of the upregulated genes in cholesterol synthesis were rapid and transient (Fig. 3C). The WT mice-specific response also included increased expression of the gene for ATP-citrate lyase (*Acly*), which is involved in the synthesis of cytosolic acetyl-CoA and oxaloacetate. Cytosolic acetyl-CoA is the building block for de novo synthesis of fatty acids and sterols. In the *ob/ob* mice, no pathway was enriched in 791 glucose-responsive genes that were specifically upregulated in this state. However, genes specifically upregulated in *ob/ob* mice included those involved in glycolysis, such as glyceraldehyde-3-phosphate dehydrogenase (*Gapdh*) and pyruvate kinase (*Pklr*), and those involved in lipid synthesis, such as glycerol-3-phosphate dehydrogenase 2 (*Gpd2*) and acetyl-CoA carboxylase beta (*Acacb*) (Fig. 3C). *Gpd1* increased in expression in *ob/ob* mice, but decreased in the WT mice (Fig. 3C, pink box). *Gpd1, Acat2*, and *Hmgcs1* are notable, because these genes exhibited different directions of regulation in the obese and healthy states.

Pathway enrichment analysis revealed a decrease in the expression of genes associated with the metabolism of linoleic acid, arachidonic acid, and vitamin A (retinol metabolism) in the WT mice (Table 1). Genes associated with ribosomes were the most significantly enriched in the downregulated genes in *ob/ob* mice (Table 1), suggesting that overall protein synthesis in the liver was reduced in response to glucose in the obese state.

To reveal the regulatory mechanism for the glucose-responsive genes, we inferred the regulatory connections between transcription factors and genes using hierarchical clustering analysis of gene expression time courses and bioinformatic analysis of binding motifs in the transcription factor database TRANSFAC (*21, 22*) (Fig. 3C, fig. S6, tables S7 and S8, and Supplementary Text). If a transcription factor binding motifs were enriched in the promoter regions of the genes in a cluster, we inferred the regulatory connections between the transcription factor and the genes in the cluster. For example, we identified the inferred regulatory connections between sterol regulatory element binding factor (Srebf) and some of the WT mice-specific upregulated genes, and between early growth response (EGR1) and some of the *ob*/*ob* mice-specific upregulated genes (Fig. 3C). We compared the inferred regulatory connections between transcription factors and genes with those predicted from chromatin immunoprecipitation (ChIP) experimental data obtained from the ChIP-Atlas database (*23*), and most of the inferred connections of the transcription factors significantly overlapped with those predicted from ChIP data (fig. S6C and table S9). The inferred regulatory connections between glucose-responsive transcription factors and glucose-responsive genes encoding metabolic enzymes served as the inter-layer regulatory connections between the TF layer and the Enzyme layer in the trans-omic network (see Fig. 5).

### Identification of glucose-responsive phosphorylation of insulin signaling molecules

Many metabolic reactions are regulated by phosphorylation either at the level of the enzyme directly through enzyme phosphorylation or at the level of the transcription factor to regulate expression of the enzyme encoding gene. Phosphorylation of enzymes and transcription factors, is regulated by signaling molecules. Glucose stimulates the release of insulin, which activates a signaling in the liver. Thus, we measured the amount of 3 proteins (insulin receptor and insulin receptor substrate 1 and 2) and the phosphorylation of 11 enzymes, transcription factors, and signaling molecules in the insulin pathway from the liver of WT and *ob*/*ob* mice following oral glucose administration (Fig. 4, fig. S8, and table S10).

**Figure 4.**
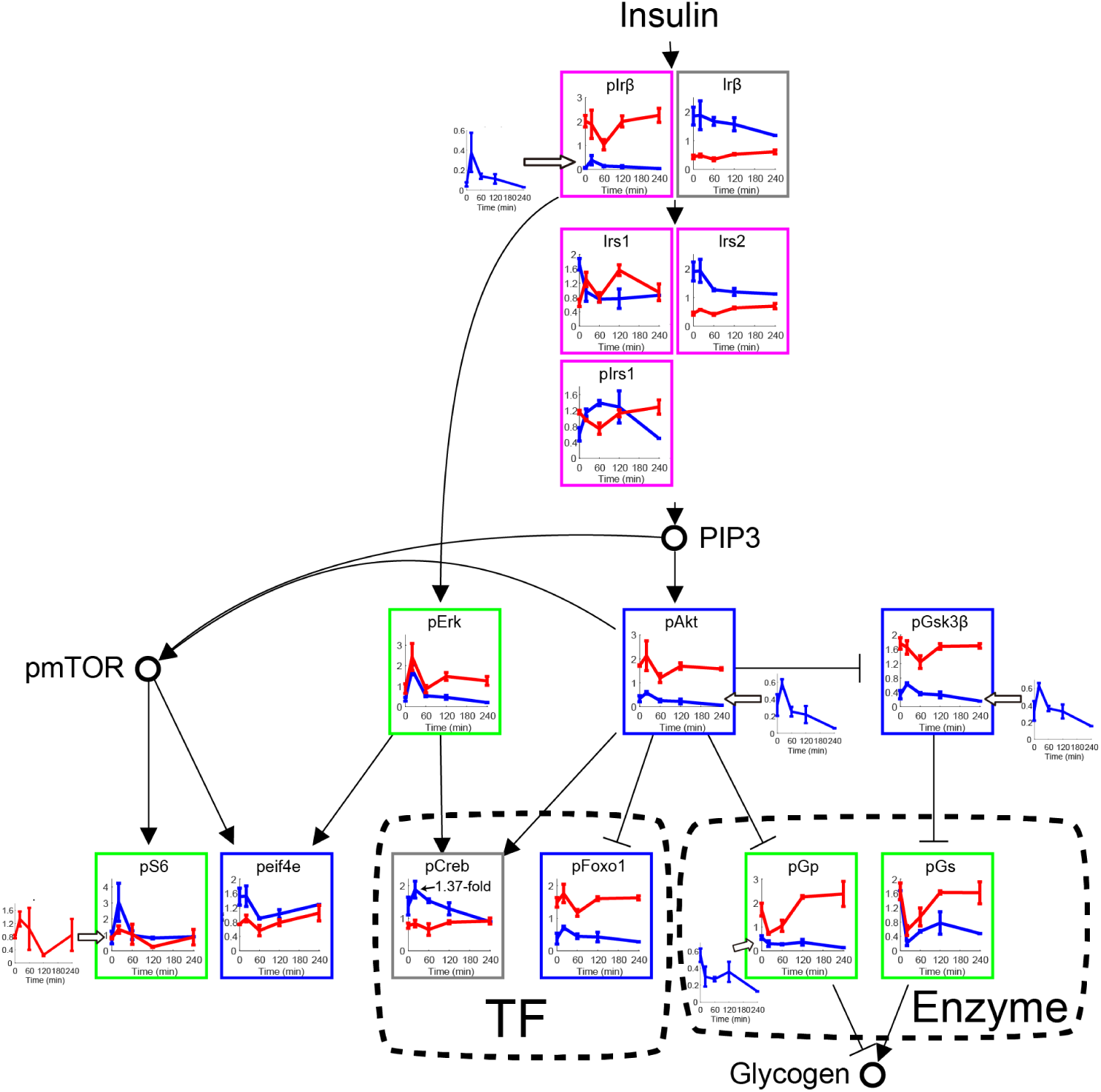
Identification of glucose-responsive phosphorylation of insulin signaling molecules. Time courses of the amount and phosphorylation of the indicated insulin signaling molecules in the liver of WT mice (blue lines) and *ob*/*ob* mice (red lines) following oral glucose administration. Phosphorylated proteins are indicated by the prefix “p” (for example, phosphorylated Erk, pErk). Data are shown as the mean and SEM of 3 mice except for WT samples for 3 time points, which had inconsistent loading (see fig. S8 for Western blot). The time course graphs are presented in the context of the insulin signaling pathway from the KEGG database (*24, 25*). Edges may reflect direct or indirect regulatory events. Not all molecules in this pathway are shown. The nodes that are presented as circles [phosphatidyl-inositol 3,4,5-trisphosphate (PIP3) and mammalian target of rapamycin (mTor)] were not quantified here. The colors of the boxes around each graph indicate a change in amount or phosphorylation specific to WT (blue), specific to *ob*/*ob* (red), common to both (green), opposite between WT mice and *ob*/*ob* (pink). Proteins that did not exhibit a change in amount or phosphorylation are outlined in gray. Proteins that showed an absolute log_2_ fold change larger than 0.585 (2^0.585^ = 1.5) at any time point were defined as glucose-responsive. Glucose-responsive molecules in the TF and Enzyme layers are enclosed in dashed boxes. See table S10 for the unabbreviated names of the insulin signaling molecules.

We identified those proteins that exhibited glucose-responsive phosphorylation that were common in liver from both WT and *ob*/*ob* mice and those that changed only in the WT or *ob/ob* mice and those that changed in opposing directions between the healthy and obese states. Changes in common were increased phosphorylation of Erk and ribosomal protein S6 and decreased phosphorylation of glycogen phosphorylase (Gp) and glycogen synthase (Gs), indicating a possible increase in protein synthesis and glycogen production. Phosphorylation of Akt, insulin receptor β (Irβ), forkhead box protein O1 (Foxo1), and glycogen synthase kinase 3 β (Gsk3β) transiently increased in WT mice but not in *ob*/*ob* mice. Several proteins exhibited changes in opposite directions between WT and *ob/ob* mice. We found that the amount of phosphorylated Irβ (pIrβ) and insulin receptor substrate 1 (pIrs1) increased in WT mice and decreased in *ob/ob* mice in response to oral glucose administration. The amounts of Irs1 and Irs2 decreased in WT mice and increased in *ob/ob* mice. These opposing responses at the early parts of the insulin pathway could contribute to the divergence in the responses, such as the increase in phosphorylated Akt only occurring in the WT mice in response to oral glucose administration. We confirmed that the phosphorylation of Akt and Erk do not increase following oral water administration (fig. S7C). Of the molecules showing glucose-responsive phosphorylation, we placed pGs and pGp in the Enzyme layer of the trans-omic network, pFoxo1 in the TF layer, and the others in the Insulin signal layer (see Fig. 5).

**Figure 5.**
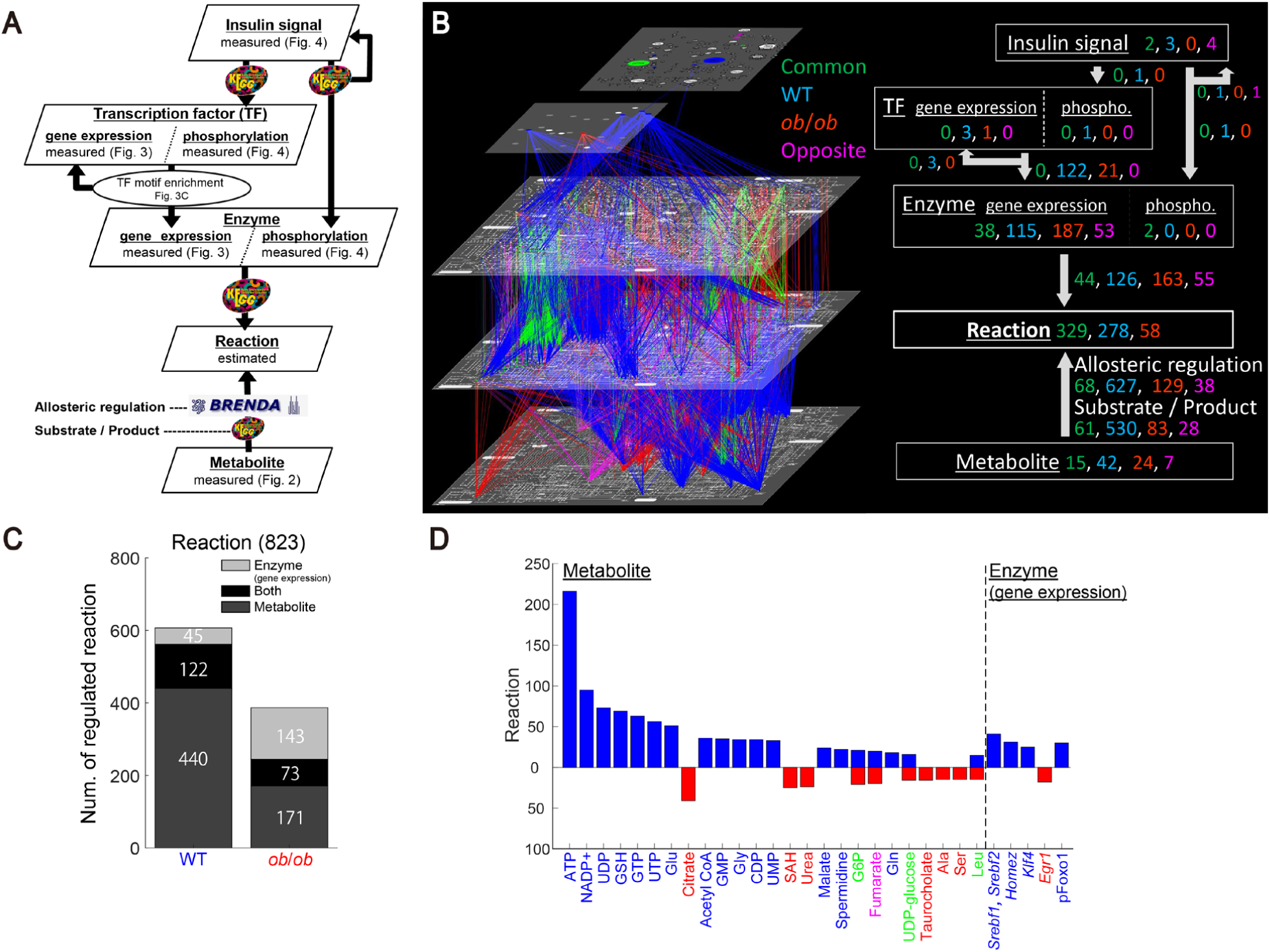
The construction of regulatory trans-omic network for glucose-responsive metabolic reactions. (**A**) The procedure for constructing the regulatory trans-omic network for glucose-responsive metabolic reactions. The layers of Insulin signal, Transcription factor (TF), Enzyme, and Metabolite correspond to glucose-responsive molecules. The Reaction layer represents “glucose-responsive metabolic reactions”, which are defined as metabolic reaction regulated by the glucose-responsive molecules. The upward and downward arrows indicate inter-layer regulatory connections, and the recurrent arrows indicate intra-layer regulatory events. The databases used to identify the inter- and intra-layer regulatory connections are shown on the arrows. (**B**) The regulatory trans-omic network for glucose-responsive metabolic reactions. The left diagram represents the network as colored nodes in the layers and edges between the layers with colored nodes representing glucose-responsive molecules and colored edges representing inter-layer regulatory connections: green, glucose-responsive molecules and inter-layer regulation common in both WT and *ob*/*ob* mice; blue, specific to WT mice; red, specific to *ob*/*ob* mice; pink, opposite responses between WT mice and *ob*/*ob* mice. A common or an opposite regulatory connection was defined if the regulating molecule of the connection is common or opposite, respectively. The numbers of each type of glucose-responsive node and edge are shown with the same colors in the network summary to the right. The Insulin signal layer is the insulin signaling pathway constructed in our previous phosphoproteomic study (*18*). The Enzyme, Reaction, and Metabolite layers are organized into global metabolic pathway (mmu01100) in the KEGG database (*24, 25*). Phospho, phosphorylation. (**C**) The number of glucose-responsive metabolic reactions regulated by glucose-responsive molecules in the enzyme layer or the metabolite layer or both from a total of 823 metabolic reactions in the liver. (**D**) The number of glucose-responsive metabolic reactions regulated by the indicated glucose-responsive molecules in WT mice (upper, blue) and *ob*/*ob* mice (lower, red). The colors of the names of molecules indicate the type of glucose-responsive molecules as described in B. Glucose-responsive metabolites that regulated more than 15 metabolic reactions and exhibited significant association with any metabolic pathway (fig. S14B) are shown. The names of the transcription factors coded by glucose-responsive genes are described in italic type, and those showing glucose-responsive phosphorylation have the prefix “p.”

### The construction of regulatory trans-omic network for glucose-responsive metabolic reactions

Using the glucose-responsive molecules from the liver analyses, we constructed the regulatory trans-omic network for glucose-responsive metabolic reactions consisting of five layers— Insulin signal, TF, Enzyme, Reaction, and Metabolite— and connections between the layers representing regulatory events (Fig. 5, A and B, and tables S11 and S12). The Insulin signal layer contains those signaling molecules showing glucose-responsive phosphorylation. The TF layer contains the “glucose-responsive transcription factors”, which were defined as the transcription factors encoded by glucose-responsive genes or those with glucose-responsive phosphorylation. The Enzyme layer contains the “glucose-responsive metabolic enzymes”, which were defined as the metabolic enzymes encoded by glucose-responsive genes or those with glucose-responsive phosphorylation. The Reaction layer contains the “glucose-responsive metabolic reactions”, which were defined as metabolic reactions regulated by the glucose-responsive metabolites, glucose-responsive metabolic enzymes, or both. For example, dephosphorylation of G6P (EC 3.1.3.9) was defined as a common glucose-responsive metabolic reaction in both WT and *ob*/*ob* mice, because dephosphorylation of G6P was regulated by a common glucose-responsive metabolite G6P and by a common glucose-responsive gene *G6pc* (fig. S1). The Metabolite layer contains the glucose-responsive metabolites. Using the information from the analysis of the liver in WT and *ob/ob* mice, we colored the nodes in each layer according to those with common changes in both WT and *ob/ob* mice and those with changes that were specific to the healthy or obese states.

We then determined inter-layer regulatory connections between glucose-responsive molecules. The inter-layer connections from the Insulin signal layer to the TF layer and to the Enzyme layer were assigned according to the regulation of transcription factors or enzymes by kinases in the KEGG database (*24, 25*). The inter-layer connections from the TF layer to the Enzyme layer were based on the inferred regulatory connections of the genes encoding metabolic enzymes by transcription factors. Note that, among all the inferred regulatory connections, only those that connect to glucose-responsive transcription factors and glucose-responsive genes are shown. The inter-layer connections from the Enzyme layer to the Reaction layer were determined by matching metabolic reactions to their corresponding metabolic enzymes according to the KEGG database (*24, 25*). The inter-layer connections from the Metabolite layer to the reaction layer consisted of two types: regulatory connections mediated by allosteric regulators and regulatory connections mediated by the substrate or product of the reaction. Allosteric regulatory connections were assigned according to the BRENDA database (*26*), and substrate or product-mediated regulatory connections were according to the KEGG database (*24, 25*). This trans-omic network included the regulatory connections for various metabolic pathways, such as glycolysis (fig. S9), glycogen metabolism (fig. S10), lipid synthesis (fig. S11), and cholesterol synthesis (fig. S12).

Comparison of the regulatory trans-omic networks for glucose-responsive metabolic reactions between WT and *ob*/*ob* mice enabled identification of WT mice-specific (Fig. 5B, blue), *ob*/*ob* mice-specific (Fig. 5B, red), and common (Fig. 5B, green) glucose-responsive molecules and inter-layer regulatory connections. Overall, the numbers of WT mice-specific glucose-responsive molecules were larger than that of *ob*/*ob* mice-specific glucose-responsive molecules except for genes expression in Enzyme layer (Fig. 5B), suggesting that the response to glucose in the obese state not only is altered but involves a reduction in the healthy response. The number of WT mice-specific inter-layer regulatory connections between the Metabolite layer and the Reaction layer was also much greater than those in the *ob/ob* mice-specific connections, suggesting that changes in metabolites dominate in the healthy response to glucose. In contrast, the *ob/ob*-specific glucose-responsive molecules in the Enzyme layer and the number of *ob/ob*-specific inter-layer connections between the Enzyme layer and the Reaction layer were greater than those of WT mice, indicating that is a major area where the response to glucose differs in the obese state.

We calculated the number of glucose-responsive metabolic reactions that were regulated by glucose-responsive metabolites in the metabolite layer, by glucose-responsive genes in the Enzyme layer, or by both in WT and *ob*/*ob* mice following oral glucose administration (Fig. 5, C and D). This analysis showed that metabolite-mediated regulation dominates the response to orally administered glucose in WT mice and that the changes in gene expression play a larger role in the response of *ob/ob* mice (Fig. 5C). The glucose-responsive metabolites specific to WT mice included the cofactors ATP and NADP+, and a large number of glucose-responsive metabolic reactions were regulated by these metabolites (216 reactions by ATP and 95 reactions by NADP+) (Fig. 5D and table S11). Because metabolites only effectively change enzyme activity when the metabolite concentration is not saturating, we quantitatively evaluated metabolite concentration compared to binding affinity, *K*_m_ and *K*_i_, for each metabolic reaction (fig. S13, table S13) to infer binding site saturation (*27*). We evaluated the saturation of ATP and NADP+ for each metabolic reaction with a binding affinity in the BRENDA database (*26*): ATP was not saturating in 41 of 81 of the glucose-responsive metabolic reactions regulated by ATP, such as the conversion of F6P to fructose 1,6-bisphosphate (F1,6P) in glycolysis, and NADP+ was not saturating in 3 of 12 of glucose-responsive metabolic reactions regulated by NADP+ (fig. S13), indicating that many of these inter-layer regulatory connections can be effective in WT mice. Indeed, together these two metabolites accounted for 50% of the regulatory connections between the Metabolite layer and the Reaction layer in WT mice and likely account for the much lower number of connections between these two layers in *ob/ob* mice. Although there were fewer inter-layer connections between the TF layer and the Enzyme layer in *ob/ob* mice, the number of metabolic enzyme-encoding genes that were specifically regulated in the *ob/ob* mice was greater than the number in the WT mice. Thus, glucose-responsive metabolites, especially coenzymes, such as ATP and NADP+, played a central role in regulation of metabolic reactions in WT mice, and glucose-responsive genes contributed a greater part of the response in *ob*/*ob* mice.

### Differences in the regulatory trans-omic networks between WT and *ob*/*ob* mice

To extract the essential differences in the regulatory networks of glucose-responsive metabolic reactions between WT and *ob*/*ob* mice, we condensed the metabolic reactions in the regulatory trans-omic networks according to the metabolic pathways through the following three procedures (Fig. 6, fig. S14, and table S14). First, we collected the related metabolic reactions in a specific metabolic pathway into one “metabolic pathway node” according to the KEGG metabolic map. Second, we selected metabolic pathway nodes that exhibited statistically significant associations with any glucose-responsive molecule (table S14) and created a “Pathway” layer with the nodes and their regulatory connections (Fig. 6A). We then classified the glucose-responsive Pathway into 3 classes: carbohydrate, lipid, and amino acid (Fig. 6, B and C). As a further simplification, we also condensed the inter-layer regulations from the Metabolite layer to the Pathway layer by removing the inter-layer connections that regulated fewer than 5 metabolic reactions, and we limited the Enzyme layer to only those metabolic enzymes encoded by glucose-responsive genes. Using this process, we made the condensed regulatory trans-omic networks for glucose-responsive reactions.

**Figure 6.**
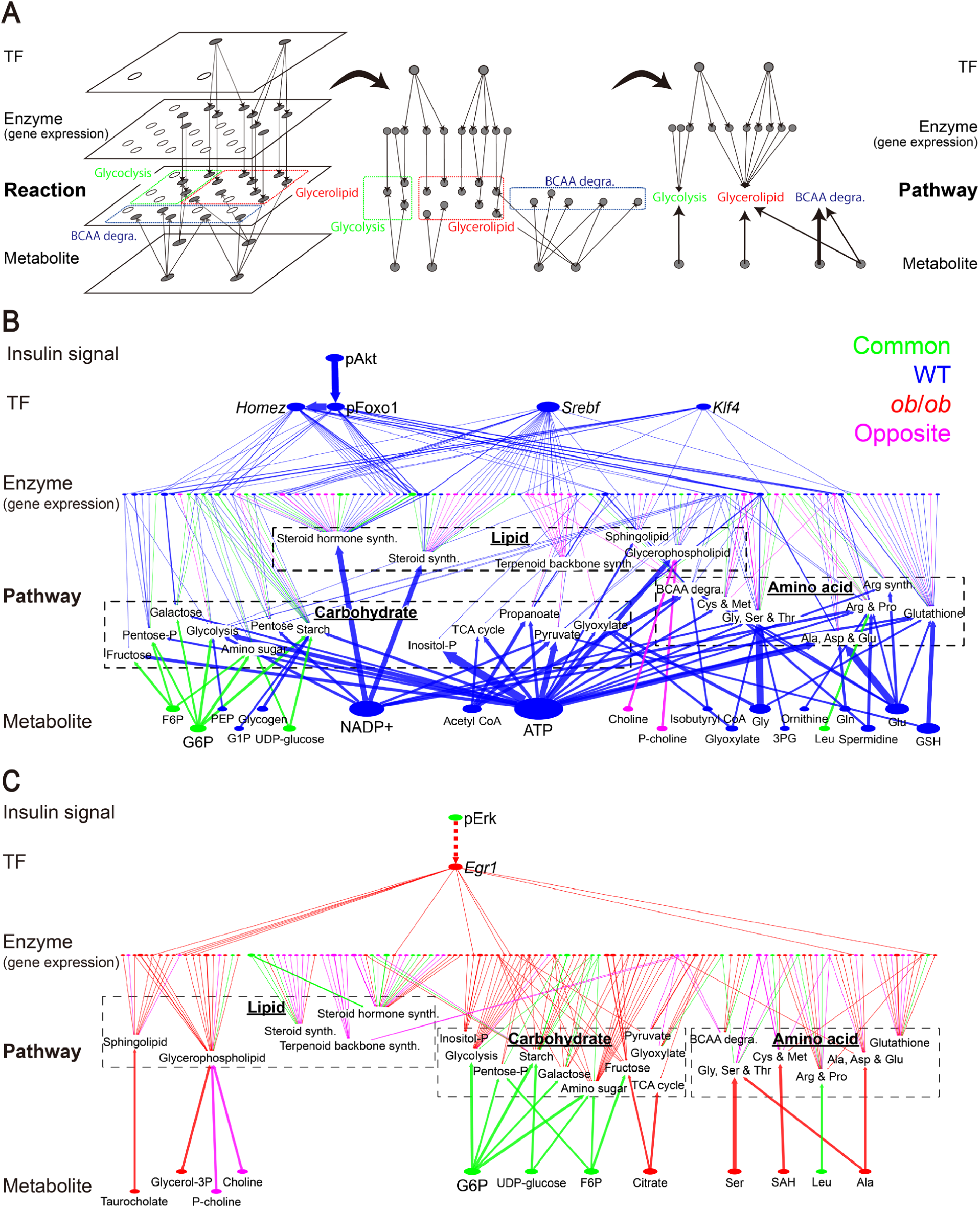
The condensed regulatory trans-omic networks for the liver metabolic response to glucose in the healthy and obese states. (**A**) The process of reducing the complexity of the trans-omic network into condensed versions. We grouped related metabolic reactions within a specific metabolic pathway into one “metabolic pathway node,” such as Glycolysis, Glycerolipid, or BCAA degradation, using the metabolic maps in KEGG (*24, 25*). We then generated a Pathway layer of the metabolic pathway nodes and grouped these nodes into classes of carbohydrate, lipid, or amino acid, according to the KEGG database (*24, 25*). (**B, C**) The condensed regulatory trans-omic network of the liver response to glucose in WT and *ob/ob* mice. The color of nodes (glucose-responsive molecules) and edges (inter-layer regulatory connections) indicates the type of molecules and regulations as described in Fig. 5B. Dashed edge between pErk and *Egr1* indicates the indirect regulatory connection (*32, 33*). In the transcription factor (TF) layer, the names of the transcription factors encoded by glucose-responsive genes are written in italics, and those showing glucose-responsive phosphorylation have the prefix “p.” *Srebf* corresponds to *Srebf1* and *Srebf2*, both of which were glucose-responsive genes specific to WT mice. The Enzyme layer contains only those metabolic enzymes that are regulated by glucose-responsive changes in gene expression, not those regulated only by phosphorylation. Metabolic pathway nodes that exhibited significant associations with any glucose-responsive molecule are included (fig. S14B). Dashed boxes enclose the nodes for the lipid, carbohydrate, and amino acid classes. Glucose-responsive metabolites that exhibited significant associations with any metabolic pathway are included. The inter-layer regulatory connections from the Metabolite layer to the Pathway layer include only those that regulate 5 or more metabolic reactions. The size of the nodes and the width of the edges indicate the relative number of the regulated metabolic reactions. See table S14 for the unabbreviated names of metabolic pathway nodes.

In the condensed regulatory trans-omic network of WT mice, we found three characteristics of the inter-layer regulatory connections that were specific to these mice (Fig. 6B): (i) from the Insulin signal layer to the TF layer and the Enzyme layer, pAkt is a major signaling molecule that directly regulates pFoxo1 and indirectly many glucose-responsive genes encoding metabolic enzymes; (ii) from the TF layer to the Enzyme layer, pFoxo1 and the glucose-responsive transcription factor-encoding genes *Srebf, Klf4*, and *Homez* regulate glucose-responsive genes encoding metabolic enzymes; and (iii) from the Metabolite layer to the Pathway layer, the glucose-responsive metabolites regulate many metabolic pathway nodes, especially in the carbohydrate and amino acid classes (see also fig. S14, A and B).

Srebf is a key transcriptional regulator of genes encoding enzymes in steroid and cholesterol biosynthesis (*28*). Indeed, the lipid class was regulated by more connections between enzymes encoded by glucose-responsive genes than was the carbohydrate class, which was regulated by more connections with metabolites (fig. S14A). Furthermore, metabolic reactions that were regulated by *Srebf*-dependent glucose-responsive genes were enriched in terpenoid backbone biosynthesis and steroid biosynthesis (fig. S14B), both of which are related to cholesterol synthesis (fig. S12). Various pathway nodes were regulated by through pFoxo1, which alleviates the transcriptional activation of its glucose-responsive target genes (fig. S14B).

In *ob*/*ob* mice, we found three characteristics of the inter-layer regulatory connections that were specific to these mice (Fig. 6C): (i) from the Insulin signal layer to the TF layer and the Enzyme layer, pErk is a signaling molecule that regulates the glucose-responsive transcription factor-encoding genes *Egr1* and glucose-responsive genes encoding metabolic enzymes downstream of this transcription factor; (ii) from the Enzyme layer to the Pathway layer, most of the regulatory connections are specific to *ob*/*ob* mice or opposing between WT mice and *ob*/*ob* mice; and (iii) from the Metabolite layer to the Pathway layer, there are few regulatory connections and the few include those common with the WT that regulate the carbohydrate class.

We considered pErk as the primary regulatory connection between the Insulin signal layer and the TF layer, because the inter-layer regulatory connection from pErk to *Egr1* and *Egr1*-regulated genes were found only in *ob*/*ob* mice. Thus, although pErk was a common glucose-responsive molecule in both WT and *ob*/*ob* mice, we assigned it to the obese regulatory network. This difference between the WT and *ob*/*ob* mice in the regulatory connections between pErk and the TF layer may relate to the sustained pErk in *ob*/*ob* mice but not in WT mice (Fig. 5) (*29, 30*). The connections between enzymes encoded by glucose-responsive genes and lipid class are more than that of carbohydrate class (fig. S14A), including the upregulated genes encoding metabolic enzymes involved in lipid synthesis, such as *Fasn* and *Gpd1* (Fig. 3B and fig. S11)

Comparing the WT and *ob*/*ob* condensed networks and their properties showed that the even these condensed networks reveal the reduction in regulatory connections in the *ob/ob* mice glucose response trans-omic network, especially between the Metabolite layer and the Pathway layer (Fig. 6, B and C, fig. S14A). Additionally, the changes in carbohydrate metabolic reactions were the most conserved between the healthy and obese states, such as the regulation of glycolysis and starch metabolism by G6P (figs. S9 and S10). The regulatory connections between the Enzyme layer and the Pathway layer were also largely different. Finally, the response was dominated by ATP and NADP+, which contributed regulatory connections to all node in the Pathway layer in WT mice (Fig. S14B).

### Differences in temporal control of the regulatory trans-omic network

We examined temporal control of glucose-responsive molecules and inter-layer regulatory connections by calculating their time constants (*T*_*1/2*_), an index of the temporal rate of response (Fig. 7 and fig. S15).

**Figure 7.**
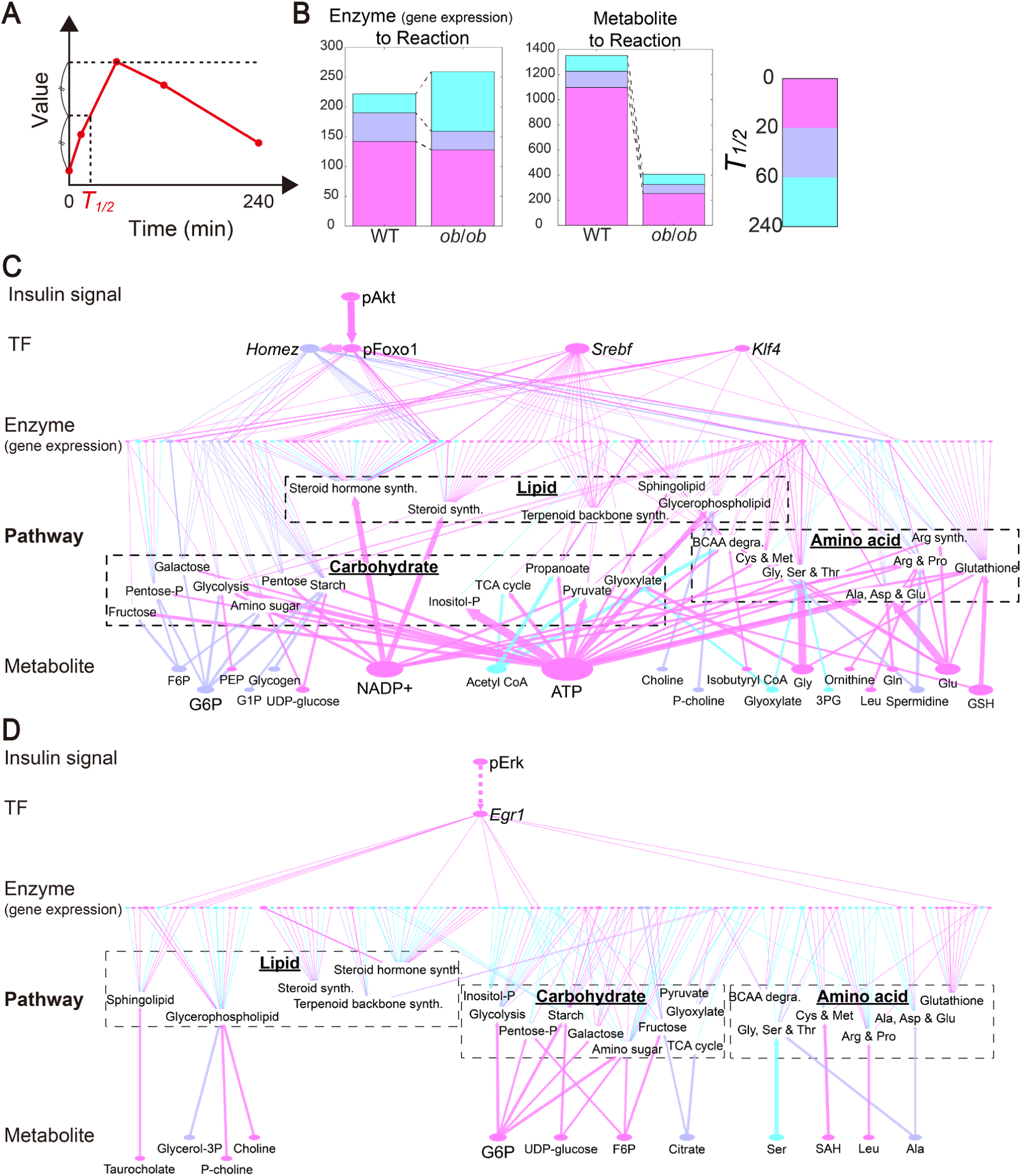
Temporal control of the regulatory trans-omic networks for the liver metabolic response to glucose in the healthy and obese states. (**A**) Definition of *T*_*1/2*_, an index of the temporal rate of response. (**B**) The number of the inter-layer regulatory connections with *T*_*1/2*_ values in the ranges indicated to the Reaction layer from the Enzyme layer and from the Metabolite layer. (**C, D**) Temporal control of the condensed regulatory trans-omic network of WT and *ob/ob* mice. The color of nodes (glucose-responsive molecules) and edges (inter-layer regulatory connections) indicates the *T*_*1/2*_ value according to the ranges in the color bar in B. *Srebf* corresponds to *Srebf1* and *Srebf2*, both of whose *T*_*1/2*_ values were 10 min. See table S14 for the unabbreviated names of metabolic pathway nodes. See table S1 for *T*_*1/2*_ values of metabolite data, table S4 for those of the enzyme- and transcription factor-encoding gene data, and table S10 for those of the phosphorylated transcription factor and signaling molecule data.

The *T*_*1/2*_ of a glucose-responsive molecule was defined as the time when the response reached the half of the maximum amplitude for an increased molecule, or of the minimum amplitude for a decreased molecule (Fig. 7A). The *T*_*1/2*_ of an inter-layer regulatory connection was defined as the *T*_*1/2*_ of a glucose-responsive molecule that is the regulating molecule of the inter-layer regulatory connection, not the *T*_*1/2*_ of the regulated molecule. Responses with *T*_*1/2*_ values shorter than 20 min were defined “rapid”, and those with values longer than 60 min were defined “slow.” From the Enzyme layer to the Reaction layer, about the half of the inter-layer regulatory connections were rapid in both WT and *ob*/*ob* mice, and the number of the slow inter-layer regulatory connections was approximately 3-times larger in *ob*/*ob* mice (Fig. 7B and fig. S15A). From the Metabolite layer to the Reaction layer, most inter-layer regulatory connections were rapid in WT mice and these were reduced in *ob*/*ob* mice.

We examined the *T*_*1/2*_ values of glucose-responsive molecules and of inter-layer regulatory connections in the condensed regulatory trans-omic networks (Fig. 7, C and D). In WT mice, most inter-layer regulatory connections were rapid (Fig. 7C). Approximately 50% of the inter-layer regulatory connections between metabolic reactions and pFoxo1-dependent glucose-responsive metabolic enzyme genes were rapid and all of the *Srebf* regulatory connections were rapid (fig. S15B). In contrast, in *ob*/*ob* mice (Fig. 7D), 50% of the inter-layer regulatory connections were slow between *Egr1*-dependent genes from the Enzyme layer to the metabolic reactions in the Pathway layer (fig. S15B). These results indicated that regulation of glucose-responsive metabolic reactions is different not only in network structure, but also in temporal control between WT mice and *ob*/*ob* mice.

## Discussion

In this study, we constructed regulatory trans-omic networks for glucose-responsive metabolic reactions in the liver of WT and *ob*/*ob* mice by integrating multi-omic data. Analysis of these networks revealed that regulation of glucose-responsive metabolic reactions in the liver was globally different between WT and *ob*/*ob* mice. In WT mice, glucose-responsive metabolic reactions were mainly regulated by the rapid response of glucose-responsive metabolites and by Akt-dependent glucose-responsive genes of metabolic enzymes. Glucose-responsive metabolites, such as coenzymes ATP and NADP+, rapidly regulated a large number of metabolic reactions in carbohydrate and amino acid metabolism, including the allosteric activation of glycogen synthesis and allosteric inhibition of glycolysis (figs. S9 and S10). A glucose-responsive gene *Srebf* was identified as the rapidly upregulated genes of metabolic enzymes related to the synthesis of cholesterol in the liver of WT mice (fig. S12), which could induce the synthesis of cholesterol esters and bile acids. This finding is consistent with the regulation of genes encoding metabolic enzymes related to cholesterol synthesis by Srebf (*31*). Because pAkt activates Srebf through mammalian target of rapamycin (mTor), and induces the expression of *Srebf* by autoregulation (*28*), the upregulation of *Srebf* might be caused by the glucose-responsive phosphorylation of Akt.

By contrast, *ob*/*ob* mice lacked most of the rapid regulation of metabolic reactions by metabolites and by *Srebf*. Instead, genes related to glycolysis, such as *Pklr* and *Gapdh*, and genes related to lipid synthesis, such as *Fasn* and *Acacb* were upregulated (figs. S9 and S10) in these obese mice. We found that *Egr1* was a glucose-responsive gene. Because we used the KEGG database to match proteins to their regulatory kinases, in our trans-omic network *Egr1* was not directly regulated by pErk. However, *Egr1* is one of the immediate early genes downstream of pErk (*32, 33*). Thus, we placed *Egr1* in the *ob*/*ob* network as regulated by pErk. In addition, the slow increase in the expression of glucose-responsive genes involved in lipid synthesis may represent a mechanism by which lipid accumulation occurs in the liver of *ob*/*ob* mice. Consistently, insulin resistance promotes de novo lipogenesis in the liver (*34*). Because lipid accumulation is thought to be one of the causes of insulin resistance (*35*), the stimulation of lipid synthesis by the upregulated genes in *ob*/*ob* mice might be one of the pathological mechanisms of lipid accumulation and insulin resistance associated with obesity. Given that the liver plays a major role of controlling postprandial blood glucose (*8, 9*), our results suggested that the rapid regulation of metabolic reactions in WT mice enables the transient increase of blood glucose following oral glucose administration, whereas the impairment of such rapid regulations and compensatory slow regulations of in *ob*/*ob* mice cause sustained hyperglycemia (fig. S2A).

The WT-specific regulation of metabolic reactions by glucose-responsive metabolites shows three unique advantages: rapidness, energetic economy, and precision. Regulation of metabolic reactions by metabolites represents a form of direct control. The metabolites serve as allosteric regulators or substrates or products, without inducing other chemical reactions or biological processes, such as protein phosphorylation and gene expression, saving the organ and organism both time and energy. The control of metabolic reactions by metabolites directly relates to the concentrations of the metabolites, enabling precise regulation of metabolic reactions. This mode of regulation contrasts with the regulation of metabolic reactions by changing the activity of the metabolic enzymes through changes in gene expression or protein phosphorylation. Controlling metabolic reactions through changes in gene expression, in particular, is less precise than controlling the reactions through changes in metabolite concentrations. Thus, our analysis indicated that the response in WT mice is faster, less energy requiring, and more precise than the response in obese mice. The application of technologies for large-scale identification of protein-metabolite interactions (*36*–*38*) will enable the validation and further analysis of the regulation of metabolic reactions by metabolites.

Our trans-omic network analysis revealed that pAkt exhibited glucose-responsive phosphorylation specifically in the liver of WT mice, whereas pErk was a common glucose-responsive event in both WT and *ob/ob* mice. Previous observations found that insulin-induced phosphorylation of Erk is not affected by obesity and that of Akt decreases in obesity (*39, 40*). Although pErk was a common glucose-responsive molecule in both WT and *ob*/*ob* mice, the inter-layer regulatory connections from pErk to *Egr1* and the *Egr1*-dependent genes were found only in *ob*/*ob* mice, but not in WT mice. The phosphorylation of Erk in *ob*/*ob* mice was sustained, whereas the phosphorylation of Erk in WT mice was transient, which could explain the difference in the role of Erk phosphorylation in the healthy and obese states. Indeed, sustained phosphorylation of Erk, but not transient phosphorylation, induces downstream gene and protein expression of immediate early genes including Egr1 (*29, 30*). Thus, the liver seems to conform to this paradigm of Erk signaling with sustained pErk representing a key property of the obese state and the slow transcriptional response to glucose administration.

Some WT-specific glucose-responsive metabolites had higher concentrations in *ob*/*ob* mice than in WT mice. In *ob*/*ob* mice, such metabolites were not glucose-responsive but were constantly maintained at higher concentrations, even during the fasting state. Thus, in the obese state these metabolic reactions are constantly influenced by such metabolites regardless of oral glucose administration. The consistently increased metabolites included ATP, some metabolites in glycolysis and in the TCA cycle and pentose phosphate pathway, and reduced glutathione (table S1), indicating constant high energy production and oxidation stress even during the fasting state in *ob*/*ob* mice. The lack of glucose-responsiveness of these metabolites in *ob*/*ob* mice may be caused by maximal energy production even during fasting state.

In this study, we focused on “glucose-responsiveness” of metabolite concentration, gene expression, and protein phosphorylation following oral glucose administration in WT and *ob*/*ob* mice. The simultaneous measurement of the time courses of multi-omic data following oral glucose administration enabled the identification of glucose-responsive molecules in each layer, and inter-layer regulatory connections. Our trans-omic networks is based on direct interaction between molecules, rather than indirect statistical relationship (*16*). The same multi-omic data enable analysis of both the difference in relative glucose-responsiveness of molecules and the difference in the amounts of molecules between WT and *ob*/*ob* mice (Supplementary Text).

Other studies have studied the difference in the amounts of molecules between healthy and obese mice using multi-omic data (*14, 41, 42*). However, none examined the difference of molecules between normal and obese mice in “glucose-responsiveness” following oral glucose administration as we did here. Soltis *et al.* (*14*) identified the epigenomic, transcriptomic, proteomic, and metabolomic differences in the liver between normal and high-fat diet (HFD)- induced obese mice during fasting state. They performed network modeling approach based on interactome and computational methods, by which they identified differences in the amounts of molecules in multi-omic network between normal and obese mice. We found similar differences by analyzing our multi-omic data from the fasting state: Obese mice had increased carbohydrate metabolism, decreased lysophospholipid metabolism, upregulation of the genes involved in carbohydrate and lipid metabolism, and downregulation of the genes involved in amino acid metabolism. Here, the *ob*/*ob* mice were fed a normal diet, not a high-fat diet, thus the common differences between the obese mice and WT mice found in both studies are likely caused by obesity, rather than by the content of diets or by an effect of lack of leptin in the *ob/ob* mice. Similar to *ob/ob* mice, diet-induced obese mice show leptin resistance and dysregulation of hepatic metabolic response (*43, 44*).

To construct the regulatory trans-omic networks for glucose-responsive metabolic reactions, we used transcriptomic and metabolomic data and data from the analysis of selected signaling proteins by Western blotting. Because of the lower comprehensiveness of data by Western blotting compared to the transcriptomic and metabolomic data, the numbers of signaling molecules, transcription factors and their inter-layer regulatory connections in our trans-omic network were poorer than those of metabolites, gene expression and their inter-layer regulatory connections. In particular, we could not identify transcription factors regulating most of the glucose-responsive genes encoding metabolic enzymes in the liver of *ob*/*ob* mice. Proteomic data will provide large-scale information about the regulation of metabolic reactions by changes in the amount of transcription factors and metabolic enzymes (*45, 46*), and phosphoproteomic data will identify the regulation of metabolic reactions by phosphorylation of transcription factors and metabolic enzymes (*15, 18, 47*). In addition, epigenomic data, as well as detailed information about the binding affinities of metabolic reactions for the metabolites that function as substrates, products, and allosteric regulators are required for constructions of a comprehensive regulatory trans-omic network for glucose-responsive metabolic reactions. Epigenomic data, including histone modification and DNA methylation, will reveal regulatory mechanisms controlling of glucose-responsive genes (*48*–*51*). Because some metabolites in the liver are regulated by transport between blood and liver, the regulation of such metabolites between blood and liver requires inclusion of the transporters that control the distribution of these metabolites (*52, 53*). Although our trans-omic network is not comprehensive, we used the network to reveal key insights into how liver metabolism is differentially regulated between the healthy and obese states in response to glucose. This in vivo study provides key insights into how obesity profoundly changes the regulatory profile, shifting regulation away from rapid regulation by metabolites to slow regulation by gene expression.

## Acknowledgments

We thank Fumio Matsuda (Osaka University) for technical advice with the analysis of metabolism and helpful discussions; Maki Ohishi, Ayano Ueno, Hiroko Maki, Keiko Endo, and Sanae Ashitani (Keio University) for their technical assistance with metabolomic analysis using CE-MS; and our laboratory members for critically reading this manuscript and for their technical assistance with the experiments. The computational analysis of this work was performed in part with support of the super computer system of National Institute of Genetics (NIG), Research Organization of Information and Systems (ROIS). This manuscript was edited by Nancy R. Gough (BioSerendipity, LLC).

## Author contributions

T.K., A. Hatano, Y.I., M.E., R.E., H.K., M.M., and K.I.N. designed and performed the animal experiments, enzymatic assays and Western blotting measurements; A. Hirayama and T.S. performed metabolomic analysis using CE-MS; K.I. and M.A. performed lipidomic analysis using LC-MS; Y.S. performed transcriptomic analysis using RNA-seq; T.K., Y.I., K.Y., S.O., M.F., K.H., and S.U. performed trans-omic analysis; writing group consisted of T.K., A. Hatano, Y.I., H.I., and S.K.; and the study was conceived and supervised by T.K. and S.K.

## Data and materials availability

Sequencing data measured in this study have been deposited in the DNA Data Bank of Japan Sequence Read Archive (DRA) (www.ddbj.nig.ac.jp/) under the accession no. DRA008416. All other data are available with the published article.

## References and Notes

1. Action to Control Cardiovascular Risk in Diabetes Study Group et al., Effects of intensive glucose lowering in type 2 diabetes. N. Engl. J. Med. 358, 2545–59 (2008).

2. Diabetes Control and Complications Trial Research Group et al., The effect of intensive treatment of diabetes on the development and progression of long-term complications in insulin-dependent diabetes mellitus. N. Engl. J. Med. 329, 977–86 (1993).

3. ADVANCE Collaborative Group et al., Intensive blood glucose control and vascular outcomes in patients with type 2 diabetes. N. Engl. J. Med. 358, 2560–72 (2008).

4. G. F. Cahill, Fuel metabolism in starvation. Annu. Rev. Nutr. 26, 1–22 (2006).

5. S. E. Kahn, R. L. Hull, K. M. Utzschneider, Mechanisms linking obesity to insulin resistance and type 2 diabetes. Nature. 444, 840–6 (2006).

6. S. V.T., S. G.I., The pathogenesis of insulin resistance: Integrating signaling pathways and substrate flux. J. Clin. Invest. 126, 12–22 (2016).

7. M. C. Petersen, D. F. Vatner, G. I. Shulman, Regulation of hepatic glucose metabolism in health and disease. Nat. Rev. Endocrinol. 13, 572–587 (2017).

8. J. Radziuk, S. Pye, Hepatic glucose uptake, gluconeogenesis and the regulation of glycogen synthesis. Diabetes. Metab. Res. Rev. 17, 250–72 (2001).

9. A. D. Cherrington, Banting Lecture 1997. Control of glucose uptake and release by the liver in vivo. Diabetes. 48, 1198–214 (1999).

10. E. A. Franzosa et al., Sequencing and beyond: Integrating molecular “omics” for microbial community profiling. Nat. Rev. Microbiol. 13, 360–372 (2015).

11. Y. Hasin, M. Seldin, A. Lusis, Multi-omics approaches to disease. Genome Biol. 18, 1–15 (2017).

12. L. Gerosa et al., Pseudo-transition Analysis Identifies the Key Regulators of Dynamic Metabolic Adaptations from Steady-State Data. Cell Syst. 1, 270–282 (2015).

13. S. R. Hackett et al., Systems-level analysis of mechanisms regulating yeast metabolic flux. Science. 354 (2016), doi:10.1126/science.aaf2786.

14. A. R. Soltis et al., Hepatic Dysfunction Caused by Consumption of a High-Fat Diet. Cell Rep. 21, 3317–3328 (2017).

15. K. Yugi et al., Reconstruction of Insulin Signal Flow from Phosphoproteome and Metabolome Data. Cell Rep. 8, 1171–1183 (2014).

16. K. Yugi, H. Kubota, A. Hatano, S. Kuroda, Trans-Omics: How To Reconstruct Biochemical Networks Across Multiple “Omic” Layers. Trends Biotechnol. 34, 276–290 (2016).

17. K. Yugi, S. Kuroda, Metabolism-Centric Trans-Omics. Cell Syst. 4, 19–20 (2017).

18. K. Kawata et al., Trans-omic Analysis Reveals Selective Responses to Induced and Basal Insulin across Signaling, Transcriptional, and Metabolic Networks. iScience. 7, 212–229 (2018).

19. P. Lindström, The physiology of obese-hyperglycemic mice [*ob*/*ob* mice]. ScientificWorldJournal. 7, 666–85 (2007).

20. R. C. Nordlie, J. D. Foster, A. J. Lange, Regulation of glucose production by the liver. Annu. Rev. Nutr. 19, 379–406 (1999).

21. V. Matys et al., TRANSFAC(R) and its module TRANSCompel(R): transcriptional gene regulation in eukaryotes. Nucl. Acids Res. 34, D108–110 (2006).

22. A. E. Kel et al., MATCH™: A tool for searching transcription factor binding sites in DNA sequences. Nucleic Acids Res. 31, 3576–3579 (2003).

23. S. Oki et al., ChIP-Atlas: a data-mining suite powered by full integration of public ChIP-seq data. EMBO Rep. 19, e46255 (2018).

24. M. Kanehisa, S. Goto, Y. Sato, M. Furumichi, M. Tanabe, KEGG for integration and interpretation of large-scale molecular data sets. Nucleic Acids Res. 40, 109–114 (2012).

25. M. Kanehisa, M. Furumichi, M. Tanabe, Y. Sato, K. Morishima, KEGG: New perspectives on genomes, pathways, diseases and drugs. Nucleic Acids Res. 45, D353–D361 (2017).

26. I. Schomburg et al., BRENDA in 2013: Integrated reactions, kinetic data, enzyme function data, improved disease classification: New options and contents in BRENDA. Nucleic Acids Res. 41, 764–772 (2013).

27. J. O. Park et al., Metabolite concentrations, fluxes and free energies imply efficient enzyme usage. Nat. Chem. Biol. 12, 482–9 (2016).

28. T. Il Jeon, T. F. Osborne, SREBPs: Metabolic integrators in physiology and metabolism. Trends Endocrinol. Metab. 23, 65–72 (2012).

29. L. O. Murphy, J. P. MacKeigan, J. Blenis, A network of immediate early gene products propagates subtle differences in mitogen-activated protein kinase signal amplitude and duration. Mol. Cell. Biol. 24, 144–53 (2004).

30. M. Ebisuya, The duration, magnitude and compartmentalization of ERK MAP kinase activity: mechanisms for providing signaling specificity. J. Cell Sci. 118, 2997–3002 (2005).

31. J. D. Horton, J. L. Goldstein, M. S. Brown, SREBPs: activators of the complete program of cholesterol and fatty acid synthesis in the liver. J. Clin. Invest. 109, 1125–31 (2002).

32. D. Gineitis, R. Treisman, Differential Usage of Signal Transduction Pathways Defines Two Types of Serum Response Factor Target Gene. J. Biol. Chem. 276, 24531–24539 (2001).

33. T. Harada, T. Morooka, S. Ogawa, E. Nishida, ERK induces p35, a neuron-specific activator of Cdk5, through induction of Egr1. Nat. Cell Biol. 3, 453–459 (2001).

34. M. Roden, Mechanisms of Disease: Hepatic steatosis in type 2 diabetes - Pathogenesis and clinical relevance. Nat. Clin. Pract. Endocrinol. Metab. 2, 335–348 (2006).

35. V. T. Samuel, G. I. Shulman, Mechanisms for insulin resistance: common threads and missing links. Cell. 148, 852–71 (2012).

36. G. X. Yang, X. Li, M. Snyder, Investigating metabolite-protein interactions: An overview of available techniques. Methods. 57, 459–466 (2012).

37. M. Diether, U. Sauer, Towards detecting regulatory protein–metabolite interactions. Curr. Opin. Microbiol. 39, 16–23 (2017).

38. I. Piazza et al., A Map of Protein-Metabolite Interactions Reveals Principles of Chemical Communication. Cell. 172, 358–372.e23 (2018).

39. Z. Y. Jiang et al., Characterization of selective resistance to insulin signaling in the vasculature of obese Zucker (fa/fa) rats. J. Clin. Invest. 104, 447–57 (1999).

40. K. Cusi et al., Insulin resistance differentially affects the PI 3-kinase- and MAP kinase-mediated signaling in human muscle. J. Clin. Invest. 105, 311–20 (2000).

41. B. W. Parks et al., Genetic architecture of insulin resistance in the mouse. Cell Metab. 21, 334–346 (2015).

42. E. G. Williams et al., Systems proteomics of liver mitochondria function. Science. 352, aad0189 (2016).

43. N. Sáinz, J. Barrenetxe, M. J. Moreno-Aliaga, J. A. Martínez, Leptin resistance and diet-induced obesity: central and peripheral actions of leptin. Metabolism. 64, 35–46 (2015).

44. P. J. Scarpace, Y. Zhang, Leptin resistance: a prediposing factor for diet-induced obesity. Am. J. Physiol. Integr. Comp. Physiol. 296, R493–R500 (2009).

45. B. Schwanhäusser et al., Global quantification of mammalian gene expression control. Nature. 473, 337–342 (2011).

46. A. R. Kristensen, J. Gsponer, L. J. Foster, Protein synthesis rate is the predominant regulator of protein expression during differentiation. Mol. Syst. Biol. 9, 689 (2013).

47. S. J. Humphrey et al., Dynamic adipocyte phosphoproteome reveals that Akt directly regulates mTORC2. Cell Metab. 17, 1009–20 (2013).

48. M. Ahrens et al., DNA methylation analysis in nonalcoholic fatty liver disease suggests distinct disease-specific and remodeling signatures after bariatric surgery. Cell Metab. 18, 296–302 (2013).

49. H. Kirchner et al., Altered DNA methylation of glycolytic and lipogenic genes in liver from obese and type 2 diabetic patients. Mol. Metab. 5, 171–183 (2016).

50. J. Lee, Y. Kim, S. Friso, S. W. Choi, Epigenetics in non-alcoholic fatty liver disease. Mol. Aspects Med. 54, 78–88 (2017).

51. C. Ling, T. Rönn, Epigenetics in Human Obesity and Type 2 Diabetes. Cell Metab. 29, 1028–1044 (2019).

52. A. Noronha et al., The Virtual Metabolic Human database: integrating human and gut microbiome metabolism with nutrition and disease. Nucleic Acids Res. 47, D614–D624 (2019).

53. M. I. Sigurdsson, N. Jamshidi, E. Steingrimsson, I. Thiele, B. T. Palsson, A detailed genome-wide reconstruction of mouse metabolism based on human Recon 1. BMC Syst. Biol. 4 (2010), doi:10.1186/1752-0509-4-140.

54. T. Soga, D. N. Heiger, Amino acid analysis by capillary electrophoresis electrospray ionization mass spectrometry. Anal. Chem. 72, 1236–41 (2000).

55. T. Soga et al., Differential metabolomics reveals ophthalmic acid as an oxidative stress biomarker indicating hepatic glutathione consumption. J. Biol. Chem. 281, 16768–16776 (2006).

56. T. Soga et al., Metabolomic profiling of anionic metabolites by capillary electrophoresis mass spectrometry. Anal. Chem. 81, 6165–74 (2009).

57. N. Ishii et al., Multiple high-throughput analyses monitor the response of E. coli to perturbations. Science. 316, 593–7 (2007).

58. K. Ikeda, in Bioactive Lipid Mediators (Springer Japan, Tokyo, 2015; http://link.springer.com/10.1007/978-4-431-55669-5_25), xpp. 349–356.

59. H. Tsugawa et al., Comprehensive identification of sphingolipid species by in silico retention time and tandem mass spectral library. J. Cheminform. 9, 19 (2017).

60. R. Noguchi et al., The selective control of glycolysis, gluconeogenesis and glycogenesis by temporal insulin patterns. Mol. Syst. Biol. 9, 664 (2013).

61. A. Von Wilamowitz-Moellendorff et al., Glucose-6-phosphate-mediated activation of liver glycogen synthase plays a key role in hepatic glycogen synthesis. Diabetes. 62, 4070–4082 (2013).

62. K. Matsumoto, A. Suzuki, H. Wakaguri, S. Sugano, Y. Suzuki, Construction of mate pair full-length cDNAs libraries and characterization of transcriptional start sites and termination sites. Nucleic Acids Res. 42 (2014), doi:10.1093/nar/gku600.

63. P. Flicek et al., Ensembl 2014. Nucleic Acids Res. 42, 749–755 (2014).

64. F. Cunningham et al., Ensembl 2015. Nucleic Acids Res. 43, D662–D669 (2015).

65. C. Trapnell et al., Differential gene and transcript expression analysis of RNA-seq experiments with TopHat and Cufflinks. Nat. Protoc. 7, 562–578 (2012).

66. C. Trapnell, L. Pachter, S. L. Salzberg, TopHat: Discovering splice junctions with RNA-Seq. Bioinformatics. 25, 1105–1111 (2009).

67. T. Sano et al., Selective control of up-regulated and down-regulated genes by temporal patterns and doses of insulin. Sci. Signal. 9, ra112 (2016).

68. C. Trapnell et al., Transcript assembly and quantification by RNA-Seq reveals unannotated transcripts and isoform switching during cell differentiation. Nat. Biotechnol. 28, 511–515 (2010).

69. J. D. Storey, A direct approach to false discovery rates. J. R. Stat. Soc. Ser. B Stat. Methodol. (2002).

70. Y. H. Yoav Benjamini, Controlling the False Discovery Rate : A Practical and Powerful Approach to Multiple Testing. J. R. Stastical Soc. 57, 289–300 (1995).

71. R. J. Kinsella et al., Ensembl BioMarts: A hub for data retrieval across taxonomic space. Database. 2011, 1–9 (2011).

72. E. Arner et al., Transcribed enhancers lead waves of coordinated transcription in transitioning mammalian cells. Science (80-.). 347, 1010–1015 (2015).

73. L. Rui, Energy metabolism in the liver. Compr. Physiol. 4, 177–97 (2014).

74. C. J. Bult et al., The Mouse Genome Database (MGD): mouse biology and model systems. Nucleic Acids Res. 36, D724–8 (2008).

75. J. Nakae, T. Kitamura, D. L. Silver, D. Accili, The forkhead transcription factor Foxo1 (Fkhr) confers insulin sensitivity onto glucose-6-phosphatase expression. J. Clin. vest, 1081359–1367 (2001).

76. A. Barthel, D. Schmoll, T. G. Unterman, FoxO proteins in insulin action and metabolism. Trends Endocrinol. Metab. 16, 183–189 (2005).

77. K. R. Albe, M. H. Butler, B. E. Wright, Cellular concentrations of enzymes and their substrates. J. Theor. Biol. 143, 163–195 (1990).

78. R. Milo, What is the total number of protein molecules per cell volume? A call to rethink some published values. BioEssays. 35, 1050–1055 (2013).

79. B. H. Junker, C. Klukas, F. Schreiber, Vanted: A system for advanced data analysis and visualization in the context of biological networks. BMC Bioinformatics. 7, 1–13 (2006).

80. G. Perriello et al., Estimation of glucose-alanine-lactate-glutamine cycles in postabsorptive humans: role of skeletal muscle. Am. J. Physiol. 269, E443–50 (1995).

81. J. Katz, J. a Tayek, Gluconeogenesis and the Cori cycle in 12-, 20-, and 40-h-fasted humans. Am. J. Physiol. 275, E537–E542 (1998).

82. M. a Herman, B. B. Kahn, Glucose transport and sensing in the maintenance of glucose homeostasis and metabolic harmony. J. Clin. Invest. 116, 1767–75 (2006).

83. O. Shaham et al., Metabolic profiling of the human response to a glucose challenge reveals distinct axes of insulin sensitivity. Mol. Syst. Biol. 4, 214 (2008).

84. Y. Shimomura et al., Branched-chain amino acid catabolism in exercise and liver disease. J. Nutr. 136, 250S–3S (2006).

85. L. Laffel, Ketone bodies: a review of physiology, pathophysiology and application of monitoring to diabetes. Diabetes. Metab. Res. Rev. 15, 412–426 (1999).

