## Supplemental Materials for "Trans-omic analysis reveals allosteric and gene regulation-axes for altered glucose-responsive liver metabolism associated with obesity"

### Materials and Methods

Mouse studies

Mouse experiments were approved by the animal ethics committee of The University of Tokyo. Ten-week-old male C57BL/6 wild-type and *ob*/*ob* mice were purchased from Japan SLC Inc. After overnight fasting (16 h), mice were administered 2 g/kg body weight of glucose or the same amount of water orally. Blood was collected from tail vein at the indicated times to measure blood glucose levels and insulin levels. At the end of the experiments, mice were sacrificed by cervical dislocation and the liver (whole or left lateral lobe) was dissected and immediately frozen in liquid nitrogen. The frozen liver was pulverized with dry ice to a fine powder with a blender and separated into tubes for omic analysis (metabolomics, lipidomics, and transcriptomics), glycogen assay, triglyceride assay, Western blotting, and quantitative real-time polymerase chain reaction (qRT-PCR). For plasma metabolomic analysis, blood was collected from retro-orbital sinus into tubes containing 0.5 mg of EDTA soon after cervical dislocation and plasma was separated by centrifugation at 2,300 × *g* for 15 min at 4ºC.

Metabolomic analysis

Total metabolites and proteins were extracted from the liver with methanol:chloroform:water (2.5:2.5:1) extraction. Approximately 40 mg of the liver was suspended with 500 µL of ice-cold methanol containing internal standards [20 µM L-methionine sulfone (Wako), 2-Morpholinoethanesulfonic acid, monohydrate (Dojindo), and D-Camphor-10-sulfonic acid (Wako)] for normalization of peak intensities of mass spectrometry (MS) among runs, then with , 500 µL of chloroform, and finally with 200 µL of water. After centrifugation at 4,600 × *g* for 15 min at 4ºC, the separated aqueous layer was filtered through a 5 kDa cutoff filter (Millipore) to remove protein contamination. The filtrate (320 µL) was lyophilized and, prior to MS analysis, dissolved in 50 µL water containing reference compounds [200 µM each of trimesate (Wako) and 3-aminopyrrolidine (Sigma-Aldrich)]. Proteins were precipitated by addition of 800 µL of ice-cold methanol to the interphase and organic layers and centrifuged at 12,000 × *g* for 15 min at 4ºC. The pellet was washed with 1 mL of ice-cold 80% (v/v) methanol and resuspended in 1 mL of sample buffer containing 1% SDS and 50 mM Tris-Cl pH8.8, followed by sonication. The total protein concentration was determined by bicinchoninic acid (BCA) assay and was used for normalization of metabolite concentration among samples.

Metabolites in plasma were extracted with methanol:chloroform:water (2.5:2.5:1). Plasma (40 µL) was extracted with sequential addition of 400 µL of ice-cold methanol containing the internal standards, 400 µL of chloroform, and 120 µL of water. After centrifugation at 10,000 × *g* for 3 min at 4ºC, the separated aqueous layer was filtered through a 5 kDa cutoff filter (Millipore) to remove protein contamination. The filtrate (300 µL) was lyophilized and, prior to analysis by MS, dissolved in 50 µL water containing the reference compounds.

All CE-MS experiments were performed using an Agilent 1600 Capillary Electrophoresis system (Agilent technologies), a G1603A Agilent CE-MS adapter kit, and a G1607A Agilent CE electrospray ionization (ESI)-MS sprayer kit. Briefly, to analyze cationic compounds, a fused silica capillary [50 µm internal Diameter (i.d.) × 100 cm] was used with 1 M formic acid as the electrolyte (*54*). Methanol/water (50% v/v) containing 0.01 µM hexakis(2,2-difluoroethoxy)phosphazene was delivered as the sheath liquid at 10 µL/min. ESI-TOFMS was performed in positive ion mode, and the capillary voltage was set to 4 kV. Automatic recalibration of each acquired spectrum was achieved using the masses of the reference standards ([$^{13}$C isotopic ion of a protonated methanol dimer (2 MeOH+H)]$^+$, *m/z* 66.0631) and ([hexakis(2,2-difluoroethoxy)phosphazene +H]$^+$, *m/z* 622.0290). To identify metabolites, relative migration times of all peaks were calculated by normalization to the reference compound 3-aminopyrrolidine. The metabolites were identified by comparing their *m/z* values and relative migration times to the metabolite standards. Quantification was performed by comparing peak areas to calibration curves generated using internal standardization techniques with methionine sulfone. The other conditions were identical to those described previously (*55*). To analyze anionic metabolites, a commercially available COSMO(+) (chemically coated with cationic polymer) capillary (50 µm i.d. x 105 cm) (Nacalai Tesque, Kyoto, Japan) was used with a 50 mM ammonium acetate solution (pH 8.5) as the electrolyte. Methanol/5 mM ammonium acetate (50% v/v) containing 0.01 µM hexakis(2,2-difluoroethoxy)phosphazene was delivered as the sheath liquid at 10 µL/min. ESI-TOFMS was performed in negative ion mode, and the capillary voltage was set to 3.5 kV. For anion analysis, trimesate and D-Camphor-10-sulfonic acid were used as the reference and the internal standards, respectively. The other conditions were identical to those described previously (*56*). We used Agilent MassHunter software (Agilent technologies) for data analysis (*55*–*57*).

<u>Lipidomic analysis</u>
Total lipids were extracted from the liver as described previously (*58*, *59*). Briefly, the frozen livers were suspended with methanol at a final concentration of 100 mg/mL of the weight of the liver. Chloroform (100 µL) and methanol (100 µL) containing internal standards [25 µM deuterated (d3-) fatty acid (FA) (16:0), 25 µM FA (18:0)-d3, 0.5 µM Cer/Sph Mixture I (Avanti Polar Lipids), 5 µM acylcarnitine (AC) (18:0)-d3, 5 µM cholic acid-d4, 0.5 µM phosphatidylglycerol 17:0/14:1, 0.5 µM lysophosphatidylcholine (LPC) (17:0), 1 µM triglyceride (24:0)- (8:0/8:0/8:0)-1,1,1-13C3] were added to 100 µL of the liver lipid mixture. After incubation for 1 h at room temperature, 20 µL of water was added and the solution was mixed and incubated for 10 min at room temperature, followed by centrifugation at 2,000 × *g* for 10 min at room temperature. The resulting supernatant was collected and injected into LC-MS systems (0.5 µL for the positive ion mode, 1 µL for the negative ion mode). LC–MS/MS analysis was performed using Triple TOF 5600+ System (SCIEX) with ACQUITY UPLC system (Waters). The reverse-phase LC separation was achieved by ACQUITY UPLC BEH column (particle size, 1.8 µm, 50 × 2.1 mm i.d., Waters Corporation) at 45°C. The mobile phase was prepared by mixing solvents (A) acetonitrile/methanol/water (20/20/60; 5 mM ammonium acetate, 10 nM EDTA) and (B) isopropanol (5 mM ammonium acetate, 10 nM EDTA) at a flow rate of 300 µL/min. Sixteen classes of lipids, each with different fatty acid composition, were analyzed, and each class was quantified by adding the area of species belonging to the class.

<u>Glycogen content assay</u>
Glycogen content was determined as previously described with some modifications (*60*). Approximately 20 mg of the liver was digested with 1.2 mL of 30% (w/v) potassium hydroxide solution for 1 h at 95°C and neutralized with 61.2 µL of glacial acetic acid. The total protein concentration of the liver digest was determined by BCA assay and adjusted to 1 µg protein/µL. Glycogen was extracted from the liver digest with Bligh and Dyer method to remove lipids (*61*). The liver digest (50 µL) was mixed with 120 µL of ice-cold methanol, 50 µL of chloroform, 10 µL of 1% (w/v) linear polyacrylamide, and 70 µL of water. After incubation on ice for 30 min,

the mixture was centrifuged at 12,000 × *g* to remove the separated aqueous layer. The glycogen was precipitated by addition of 200 µL of methanol and centrifugation at 12,000 × *g* for 30 min at 4ºC, washed with ice-cold 80% (v/v) methanol and dried completely. The glycogen pellets were suspended in 20 µL of 0.1 mg/mL amyloglucosidase (Sigma-Aldrich) in 50 mM sodium acetate buffer and incubated for 2 h at 55ºC to digest glycogen. The concentration of the glucose produced from the glycogen was determined using the Amplex Red Glucose/Glucose Oxidase Assay kit glucose assay (Thermo Fisher Scientific), according to manufacturer's instruction.

Triglyceride assay

Total triglyceride was extracted with methanol:chloroform:water (2.5:2.5:1). Livers were suspended with methanol at final concentrations of 100 mg/mL (WT) or 25 mg/mL (*ob*/*ob*) of the weight of the liver. The suspension (800 µL) was transferred into a new tube and mixed with chloroform (800 µL) and water (320 µL), followed by centrifugation at 4,600 ×*g* for 10 min at 4 ºC. The organic phase (400 µL) was collected into a new tube and resuspended in 20 µL of Triton X-100. Samples were dried and resuspended in 180 µL water. The concentration of triglyceride in the samples was determined using Triglyceride E test (Wako), according to the manufacturer's instruction.

RNA sequencing

Total RNA was extracted from the liver using RNeasy Mini Kit (QIAGEN) and QIAshredder (QIAGEN) and assessed for quantity using Nanodrop (Thermo Fisher Scientific) and for quality using the 2100 Bioanalyzer (Agilent Technologies). cDNA libraries were prepared using SureSelect strand-specific RNA library preparation kit (Agilent Technologies). The resulting cDNAs were subjected to 100-bp paired-end sequencing on an Illumina HiSeq2500 Platform (Illumina) (*62*).

Sequences were aligned to the mouse reference genome obtained from Ensembl database (*63*, *64*) (GRCm38/mm10, Ensembl release 70) using the software package TopHat (v.2.0.9) (*65*, *66*), software in the Tuxedo tool. Cufflinks (v.2.2.1), software in the Tuxedo tool, was used to assemble transcript models from aligned sequences and to estimate the number of transcripts as an indication of gene expression (*65*, *66*). The number of transcripts was shown as fragments per kilobase of exon per million mapped fragments (FPKM).

qRT-PCR

qRT-PCR was performed as previously described (*67*). Briefly, total RNA was extracted from the liver using RNeasy Mini Kit (QIAGEN) and QIAshredder (QIAGEN) and reverse-transcribed into complementary DNA (cDNA) using the High-Capacity RNA-to-cDNA Kit (Applied Biosystems), according to the manufacturer's instructions. The cDNA samples were amplified using Power SYBR Green PCR Master Mix (Applied Biosystems) and Step ONE plus Real-Time PCR system (Applied Biosystems), according to the manufacturer's protocols. The primer sequences used in the qRT-PCR analysis are listed in table S6. The fold change was calculated using the mean expression of *Rpl4* and *Gusb* as reference.

Western blotting

Total proteins were extracted from the liver with methanol:chloroform:water (2.5:2.5:1) extraction. Ice-cold methanol was added to the liver at a concentration of 100 mg/mL of the weight of the liver, and the suspension (400 μL) was mixed with chloroform (400 μL) and water (160 μL), followed by centrifugation at 4,600 × *g* for 10 min at 4ºC. The aqueous and organic phases were removed and 800 μL of ice-cold methanol was added to the interphase to precipitate proteins. The resulting pellet was suspended with 400 μL of lysis buffer [10 mM Tris-HCl (pH 6.8) in 1% SDS] and incubated for 15 min at 65ºC, followed by sonication. The protein lysate was centrifuged 12,000 × *g* for 3 min at 4ºC to remove debris. The total protein concentration of the resulting supernatant was determined by BCA assay. Antibodies were purchased from Cell Signaling Technology, as follows: pIrβ (Tyr1150/Tyr1151) (#3024), pErk1/2 (Thr202/Tyr204) (#9101), phosphorylated cAMP responsive element binding protein (pCreb) (Ser133) (#9198), phosphorylated eukaryotic translation initiation factor 4e (peif4e) (Ser209) (#9741), pAkt (Ser473) (#9271), pS6 (Ser235/Ser236) (#2211), pGsk3β (Ser9) (#9336), pGs (Ser641) (#3891), pFoxO1 (Ser256) (#9461), and β-Actin (#4967). Antibodies against total Irβ (sc-711) and β-Actin (sc-47778) were from Santa Cruz Biotechnology. pGp (Ser15) was made in house as previously described (*60*). The proteins (2 μg or 10 μg) were separated on SDS-PAGE and blotted with the antibodies. Immunodetection was performed using Immobilon Western Chemiluminescent HRP Substrate (Millipore) or SuperSignal West Pico PLUS Chemiluminescent Substrate (Thermo Fisher Scientific), and the Western blot signals were detected using a luminoimage analyzer (LAS-4000; Fujifilm or ChemiDoc Touch MP; Bio-Rad) and quantified with the TotalLab TL120 analysis software (Nonlinear Dynamics). β-Actin was used as a loading control.

Identification of glucose-responsive molecules

Molecules that were detected in less than half of replicates in either WT mice or *ob*/*ob* mice at any time point after oral glucose administration were removed from the analysis. A molecule with a statistically significant change in response to oral glucose administration was defined as a glucose-responsive molecule according to the following criteria. The fold change of the mean amount at each time point over the mean amount at fasting state (0 min) was calculated for each molecule. The significance of the change at each time point was tested by two-tailed Welch's *t*-test for each metabolite, and by CuffDiff (v. 2.2.1), software in the Tuxedo tool (*65*, *68*), for each gene. Metabolites and genes that showed an absolute $\log_2$ fold change larger than 0.585 ($2^{0.585}$ = 1.5) and an FDR-adjusted p value (q value) less than 0.1 at any time point were defined as glucose responsive metabolites (Fig. 2) and genes (Fig. 3). The q values were calculated by Storey's procedure (*69*). For Western blotting data, proteins that showed an absolute $\log_2$ fold change larger than 0.585 at any time point, except for 240 min of WT mice, were defined as glucose-responsive (Fig. 4). Because of the small number of WT mice at 240 min (n = 1), we did not use the fold changes at the time point. To define an increase or decrease in time courses with changes in both directions at different times, we used the direction of change compared to time 0 at the earliest time point that showed a significant change. Metabolites that responded to oral water administration were determined by the same procedure as the identification of glucose-responsive metabolites.

Identification of differences of the amounts of molecules at 0 min between WT mice and *ob*/*ob* mice

The differences of the amounts of molecules between WT mice and *ob*/*ob* mice before glucose administration (0 min) were determined by fold change and statistical test using the following procedure. The fold change of the mean amount of *ob*/*ob* mice at 0 min over the mean amount of

WT mice at 0 min was calculated for each molecule. The significance of differences were tested by two-tailed Welch's *t*-test for each metabolite and phosphorylation, and by CuffDiff (v. 2.2.1) (*65*, *68*) for each gene expression. Molecules that showed an absolute $\log_2$ fold change larger than 0.585 ($2^{0.585} = 1.5$) and a q value less than 0.1 were defined as different molecules in amount between WT mice and *ob*/*ob* mice. The q values were calculated by Storey's procedure for polar metabolite and gene expression (*69*). The q values were calculated by Benjamini–Hochberg procedure for lipid and phosphorylation (*70*), because of the small numbers of molecules.

Clustering analysis

Time courses for each metabolite of WT mice and *ob*/*ob* mice were normalized by dividing by the geometric mean of the values of WT mice and *ob*/*ob* mice in fasting state (0 min) and then $\log_2$-transformed. We combined the two time courses of WT and *ob*/*ob* mice for each metabolite and performed hierarchical clustering of the combined time courses using Euclidean distance and Ward's method (Fig. 2, fig. S3). Based on the clustering tree, we defined eight different clusters of metabolites, showing similar or different responses between WT mice and *ob*/*ob* mice.

Time courses for the expression of each gene of WT mice and *ob*/*ob* mice were normalized by subtracting the average of the expression values of time courses of both mice and then dividing the resulting values by the standard deviation (Z-score normalization). We combined the 2 time courses of WT and *ob*/*ob* mice for each gene and performed hierarchical clustering of the combined time courses using Euclidean distance and Ward's method (Fig. 3, fig. S6). The genes with significant differences between WT mice and *ob*/*ob* mice at any time point (q value $< 0.1$) or a significant response at any time point either in WT mice or *ob*/*ob* mice (q value $< 0.1$) were selected for the clustering analysis (7845 genes). For the selection, p value was calculated by CuffDiff (v. 2.2.1) (*65*, *68*), and q value was calculated by Storey's procedure (*69*).

Pathway enrichment analysis

We performed pathway enrichment analysis of glucose-responsive genes (Table 1 and table S5), genes showing the differences in the amounts of expression between WT mice and *ob*/*ob* mice before oral glucose administration (table S5), and the genes in each cluster (fig. S6). The enrichment of the genes in each pathway was determined using one-tailed Fisher's exact test. We used the genes detected in more than half of the replicates in WT mice and *ob*/*ob* mice at all time points as a background. In pathway enrichment analysis of the glucose-responsive genes (Table 1), we used the pathways in Metabolism, Genetic Information Processing, and Cellular Processes from the KEGG database (*24*, *25*). We used the same pathways in enrichment analysis of the genes showing the differences in the amounts of expression between WT mice and *ob*/*ob* mice before oral glucose administration. In pathway enrichment analysis of the genes in each cluster (fig. S6), we used pathways in Metabolism and Genetic Information Processing from the KEGG database (*24*, *25*). We also used the KEGG pathway "cholesterol metabolism", which included terpenoid backbone biosynthesis (mmu00900) and steroid biosynthesis (mmu00100).

Prediction of transcription factor binding motif and inference of regulatory connections between TFs and genes

The flanking regions around the major transcription start site of genes were extracted from GRCm38/mm10 (Ensembl, release 71) using Ensembl BioMart (*71*). The region from -300 bp to +100 bp of the major transcription start site was defined as the flanking region, according to the

FANTOM5 analysis of time course (*72*). The transcription factor binding motifs in each flanking region (fig. S6B) were predicted using TRANSFAC Pro, a transcription factor database, and Match, a transcription factor binding motifs prediction tool (*21*, *22*). The threshold for each transcription factor binding motif prediction was set using extended vertebrate_non_redundant_min_FP.prf, a parameter set in TRANSFAC Pro. Because some of the transcription factors known to regulate the metabolism of liver (*73*) are not included in this parameter set, we extracted the transcription factor binding motifs of PPARA, PPARG, FOXO1, and CHREBP1 from vertebrate_non_redundant.prf, and appended these transcription factor binding motifs and their parameters to vertebrate_non_redundant_min_FP.prf. If a transcription factor has multiple binding motifs, we selected the binding motif that was based on the largest number of sequences.

For the inference of regulatory connections between TFs and genes, we performed transcription factor motif enrichment analysis of the genes in each cluster (fig. S6B). The enrichment of transcription factor binding motif in the flanking regions of genes in each cluster was determined by one-tailed Fisher's exact test, and transcription factor binding motifs with q value less than 0.1 were defined as significantly enriched. The q values were calculated by Benjamini–Hochberg procedure (*70*). We used the genes analyzed in the hierarchical clustering as a background. To reduce the number of statistical tests, the clusters that contain 100 genes or more were analyzed. If a transcription factor binding motifs were enriched in the promoter regions of the genes in a cluster, we inferred the regulatory connections between the corresponding transcription factor and the genes in the cluster. To avoid overestimation, we examined the enrichment of transcription factor binding motif in two children clusters of a cluster, and excluded the parent cluster from the inference if the transcription factor binding motif was significantly enriched in one of the two children clusters. The enrichment of transcription factor binding motif in the flanking regions of genes in each of two children clusters was determined by one-tailed Fisher's exact test, and transcription factor binding motifs with p value less than 0.01 were defined as significantly enriched. We used the genes in the parent cluster as a background.

For the validation of the inferred regulatory connections, we examined the overlap between the inferred genes of each transcription factor and those predicted from experimental ChIP data from the ChIP-Atlas database (*23*) (fig. S6C). The genes for which ChIP-seq peaks of a transcription factor were detected in the flanking region around the transcription start sites were obtained using "Target genes," a prediction tool in ChIP-Atlas. We used the flanking regions from -1000 bp to +1000 bp of the transcription start sites in Target Genes. The overlap between the inferred genes and genes from ChIP data was determined by one-tailed Fisher's exact test, and those with p value less than 0.01 were defined as significant.

<u>Insulin signaling pathway</u>

Insulin signaling pathway in Fig. 4 is a subset of the nodes of the insulin signaling pathway in the KEGG database (mmu04910) (*24*, *25*) with regulatory input to cAMP responsive element binding protein (Creb) from the PI3K-Akt signaling pathway (mmu04151) and MAPK signaling pathway (mmu04010) in the KEGG database.

<u>Construction of regulatory trans-omic network of glucose-responsive metabolic reaction</u>

The regulatory trans-omic networks for glucose-responsive metabolic reactions consisted of five layers— Insulin signal, TF, Enzyme, Reaction, and Metabolite— with inter- and intra-layer regulatory connections (Fig. 5). The Insulin signal layer is the insulin signaling pathway constructed in our previous phosphoproteomic study (*18*). We included in the Insulin signal layer signaling molecules that we analyzed by Western blotting for abundance or phosphorylation state or both; we did not include transcription factors, such as Foxo1, or metabolic enzymes, such as Gs, in this layer. The TF layer consisted of all transcription factors with the inferred regulatory connection (fig. S6). The Enzyme layer consisted of all metabolic enzymes in the pathways in Metabolism obtained from the KEGG database (*24*, *25*). The Reaction layer consisted of the metabolic reactions (based on EC number) corresponding to the metabolic enzymes in the Enzyme layer. The Metabolite layer consisted of all metabolites analyzed by CE-MS. Only the molecules and reactions corresponding to the genes that were expressed in at least one sample were included in the Insulin signal, TF, Enzyme, and Reaction layers. Not all 14,292 genes were included in the network.

Each layer included the corresponding glucose-responsive molecules. The Insulin signal layer involved signaling molecules showing glucose-responsive phosphorylation (Fig. 4). The TF layer included "glucose-responsive transcription factors," which were defined as the transcription factors encoded by glucose-responsive genes or those showing glucose-responsive phosphorylation (Figs. 3 and 4). To avoid overestimation, the transcription factors with downstream genes that were not enriched in glucose-responsive upregulated genes or downregulated genes were excluded from glucose-responsive transcription factors. We used the inferred regulatory connections for the identification of upstream of glucose-responsive genes (fig. S6). The enrichment of the downstream genes in glucose-responsive genes was determined by one-tailed Fisher's exact test, and an enrichment with a q value less than 0.1 was defined as significant (table S12). The q values were calculated by Benjamini–Hochberg procedure (*70*).

The Enzyme layer included "glucose-responsive metabolic enzymes," which were defined as the metabolic enzymes encoded by glucose-responsive genes or those showing glucose-responsive phosphorylation (Figs. 3 and 4). The Reaction layer included "glucose-responsive metabolic reactions," which were defined as metabolic reactions regulated by the glucose-responsive metabolites or the glucose-responsive metabolic enzymes or both. The Metabolite layer included the glucose-responsive metabolites (Fig. 2).

We also determined the direction of glucose-responsiveness, increase or decrease. To determine a direction for time courses with both increased and decreased time points, we used the direction of change at the earliest time point with a significant difference from time 0 (fasting state). We did not determine a direction (increase or decrease) for metabolic reactions, because we did not measure metabolic reaction activity.

To determine inter-layer and intra-layer regulatory connections, we used a process we have used previously (*24*–*26*). To determine regulatory connections from the Enzyme and Metabolite layers to Reaction layer, both the target of the regulatory connection (a metabolic reaction) and the regulating molecule (enzyme or metabolite) had to be glucose responsive. Among the Insulin signal, TF, and Enzyme layers, the inter-layer regulatory connections and intra-layer regulatory connections were determined using the directions of glucose-responsiveness of the regulating molecule and the regulated molecules, and the types of the inter-layer and intra-layer regulatory connections, which were either designated positive or negative. We defined the positive inter-layer and intra-layer regulatory connections when both the regulating molecule and regulated molecule showed the same direction of change, that is both increased or both decreased. We defined the negative inter-layer and intra-layer regulatory connections when the regulating molecule and regulated molecule showed responses in the opposite direction, that is one increased and the other decreased.

The inter-layer regulatory connections from the Insulin signal layer to the TF layer and the Enzyme layer and the intra-layer regulatory connections in the Insulin signal layer mediated by phosphorylation were determined based on the kinase-substrate relationship in the insulin signaling pathway constructed in our previous phosphoproteomic study (*18*). The insulin signaling pathway is a pathway comprise of several signaling pathways in the KEGG database (*24*, *25*). The inter-layer regulatory connections from the TF layer to the Enzyme layer and the intra-layer regulatory connections in the TF layer were determined based on the relationship between transcription factors and gene expression inferred from transcription factor binding motif enrichment analysis. The types of the regulatory connections made by glucose-responsive transcription factors were defined according to the Gene Ontology Annotations obtained from the Mouse Genome Database (*74*) (table S8). The transcription factors that were included in the list of DNA-binding transcription repressor (GO:0001227) and not in the list of DNA-binding transcription activator (GO:0001228) were defined as the transcription repressors. Foxo1 was added to the list of transcription activator based on the previous studies of gluconeogenesis (*75*, *76*). The effects of the phosphorylation of transcription factors on the types of regulatory connections were defined according to the KEGG database (*24*, *25*). The inter-layer regulatory connections from the Enzyme layer to the Reaction layer were determined by regulation of metabolic reactions by the corresponding metabolic enzymes according to the KEGG database (*24*, *25*). The inter-layer regulatory connections from the Metabolite layer to the Reaction layer were of two types: regulation by allosteric regulator and regulation by the reaction’s substrate or product. Allosteric regulatory connections were determined by allosteric regulation of metabolic reactions by metabolites according to the BRENDA database (*26*). We used the allosteric regulation reported for mammals (Bos taurus, Felis catus, Homo sapiens, “Macaca,” “Mammalia,” “Monkey,” Mus booduga, Mus musculus, Rattus norvegicus, Rattus rattus, Rattus sp., Sus scrofa, “dolphin,” and “hamster”). Substrate and product regulatory connections were determined by regulation of metabolic reaction by its substrate or product according to the KEGG database (*24*, *25*). Because the reversibility of metabolic reactions was not determined, metabolic reactions were assumed to be regulated by both the substrate and product.

In the comparison of the trans-omic networks between WT mice and *ob*/*ob* mice, we identified 4 types of regulatory connections; WT mice-specific, *ob*/*ob* mice-specific, common, and opposite regulatory connections. The regulatory connections were defined as WT mice-specific or *ob*/*ob* mice-specific if the connections were identified only in WT mice or *ob*/*ob* mice, as common if the connections were identified in both WT mice and *ob*/*ob* mice and the regulating molecules were common glucose-responsive molecules, as opposite if the connections were identified in both WT mice and *ob*/*ob* mice and the regulating molecules showed opposite responses.

<u>Generation of a condensed trans-omic network based on metabolic pathway information</u>

We condensed the regulatory trans-omic networks according to the following procedures (Fig. 6). First, we grouped the related metabolic reactions in a specific metabolic pathway into one “metabolic pathway node,” according to the KEGG metabolic map (*24*, *25*). Second, we selected metabolic pathway nodes that exhibited significant associations with any glucose-responsive metabolites or transcription factors and created a “Pathway” layer with the nodes and their regulatory connections. We then classified the metabolic pathway nodes into 3 classes —

carbohydrate, lipid, and amino acid— according to the KEGG database (*24*, *25*). The association between the metabolic reactions in a metabolic pathway and those regulated by a glucose-responsive molecule was determined by one-tailed Fisher's exact test, and associations with a p value less than 0.05 were defined as significant. We also selected glucose-responsive metabolites that exhibited significant associations with any metabolic pathway nodes. Third, we reduced the inter-layer regulatory connections from the Metabolite layer to the Pathway layer by removing the inter-layer regulatory connections that regulated fewer than five metabolic reactions. We connected *Egr1* to pErk as an indirect downstream transcription factor on the previous observation (*32*, *33*) (dashed arrow).

Determination of the values of $K_m$ and $K_i$ for metabolic enzymes

We extracted $K_m$ values of a substrate for metabolic enzyme, and $K_i$ value of a metabolite for metabolic enzyme from *Mus musculus* and *Homo sapiens* from the BRENDA database (*26*) (fig. S13). The values for mutant enzymes were excluded. We used the values for *Mus musculus* unless they were not available, then we used the values for *Homo sapiens*. When multiple values for the same enzyme and metabolite were present, we used their geometric mean. To compare with the metabolomic data, we changed the unit of the metabolomic data using the weight of protein per volume in the mouse liver (0.2 mg/L) (*77*, *78*).

Implementation

Statistical tests, clustering analysis, enrichment analysis, and trans-omic network analysis were done using MATLAB 2017a (The Mathworks Inc.). Visualization of trans-omic network in Graph Modeling Language (GML) formats was done using Python 2.7 and VANTED (*79*).

### Supplementary Text

The differences of the amounts of molecules between WT mice and *ob*/*ob* mice before oral glucose administration (Figs. 2 to 4, and fig. S4)

We identified the significant differences of the amounts of metabolites, gene expression, and signaling molecules between WT mice and *ob*/*ob* mice before oral glucose administration (tables S1, S2, S4, S5, and S10). Forty six metabolites (28% of the total quantified polar metabolites) showed higher concentrations in *ob*/*ob* mice than in WT mice, including ATP, G6P, PEP, citrate, 6-phosphogluconate, phosphoenolpyruvate (PEP), sedoheptulose-7-phosphate (S7P) and reduced glutathione (table S1). Twenty one metabolites (13%) showed higher concentrations in WT mice than in *ob*/*ob* mice, most of which were non-proteinogenic amino acids. Triglyceride (TG) and acylcarnitine (AC) showed higher concentrations in *ob*/*ob* mice than in WT mice, and 8 lipids showed higher concentrations in WT mice than in *ob*/*ob* mice, including lysophosphatidylcholine (LPC) and fatty acid (FA) (table S2).

One thousand four hundred seventy four genes (10% of the total quantified genes) showed higher expressions in *ob*/*ob* mice than in WT mice (table S4), and these were enriched in glycolysis, fatty acid synthesis, and cholesterol synthesis ($p < 0.01$, table S5). Seven hundred eighty genes (5%) showed higher expressions in WT mice than in *ob*/*ob* mice, and these were enriched in six amino acid metabolic pathways ($p < 0.01$). Irβ, Akt, Foxo1, Gsk3β, and Gp showed significantly higher phosphorylation in *ob*/*ob* mice than WT mice (table S10), consistent with the higher blood insulin concentration in *ob*/*ob* mice (fig. S2A). The amount of Irβ was lower in *ob*/*ob* mice than in WT mice.

Correlation between the response of metabolite in the liver and that in the blood (fig. S5)
Some metabolites are regulated not only in the liver but also in the blood through systemic circulation. These contribute to glucose homeostasis. For example, some lactate and alanine produced in muscle is transported in the circulation, from which it is taken up into liver and converted into glucose through gluconeogenesis. These processes are the Cori cycle (lactate) and the glucose-alanine cycle (*80–82*). Therefore, we analyzed the correlation of the same set of metabolites in liver and blood. We measured the time courses of 108 metabolites in the blood of WT and *ob*/*ob* mice following oral glucose administration using CE-MS (fig. S5A and table S3). For each metabolite that was measurable in both liver and blood (86 metabolites), we calculated the correlation between the time course of the metabolite in the liver and that in the blood (fig. S5B). In both WT and *ob*/*ob* mice, branched chain amino acids (BCAAs) and 3-OH-butyrate had positive correlations between liver and blood (fig. S5C). The decrease of these metabolite in the blood is consistent with the responses in human blood following oral glucose administration (*83*). Because the activity of branched-chain aminotransferase (Bcat) is much lower in the liver than in the muscle, the decrease of BCAAs in the liver may be indirectly regulated through the blood (*84*). In contrast, because ketone bodies are mainly produced in the liver (*85*), the decrease of the ketone body 3-OH-butyrate in the blood may reflect the regulation of ketogenesis in the liver.

Hierarchical clustering of time courses of transcriptomic data (fig. S6)
We performed hierarchical clustering of the time courses of gene expression in the liver of WT and *ob/ob* mice, and pathway and transcription factor motif enrichment analysis of genes in each cluster (fig. S6 and table S7). Gene expression was divided into three main clusters characterized by the difference of expression between WT and *ob*/*ob* mice: The expression of genes in cluster 3 (3329 genes, 23% of the quantified genes) were higher in WT mice than in *ob*/*ob* mice; those in cluster 4 (3028 genes, 21%) were higher in *ob*/*ob* mice than in WT mice; those in cluster 6 (1488 metabolites, 10%) showed no differences in the amounts of expression between WT mice and *ob*/*ob* mice. Genes in cluster 3 were significantly enriched for the Foxo1 binding motif. Nuclear localization of Foxo1 is inhibited by insulin-induced phosphorylation (*75*, *76*). Therefore, the enrichment of the Foxo1 binding motif in the highly expressed genes in WT mice is consistent with the higher phosphorylation of Foxo1 in the liver of *ob*/*ob* mice (Fig. 4). Genes in cluster 3 were also enriched genes related to mRNA translation. Approximately 40% of ribosomal protein-encoding genes showed significantly higher expression in WT mice than in *ob*/*ob* mice.

Cluster 3 contained cluster 5 (2344 genes, 16%), which showed a difference in the expression between WT and *ob*/*ob* mice at time 0, and cluster 12 (985 genes, 7%), which showed the difference in the response following glucose administration between the mice. The average expression of genes in cluster 5 was higher in WT mice than in *ob*/*ob* mice following glucose administration. This cluster exhibited significant enrichment of genes encoding enzymes involved in amino acid metabolism, consistent with the higher concentration of non-proteinogenic amino acids in WT mice than in *ob*/*ob* mice (Supplementary text: The differences of the amounts of molecules between WT mice and *ob*/*ob* mice before oral glucose administration). This cluster included the genes encoding transaminases, such as glutamic-oxaloacetic transaminase 1 (*Got1*) and glutamic pyruvic transaminase (*Gpt*), the genes encoding enzymes in urea circuits, such as argininosuccinate synthetase 1 (*Ass1*), argininosuccinate lyase (*Asl*), arginase (*Arg1*), and ornithine transcarbamylase (*Otc*), and gluconeogenic genes, such as phosphoenolpyruvate carboxykinase 1 (*Pck1*) and fructose bisphosphatase 1 (*Fbp1*). *Pck1* expression is regulated by Foxo1 (*75*, *76*) . The average expression of genes in cluster 12 showed transient increases in WT mice and sustained decreases in *ob*/*ob* mice after glucose administration. This cluster showed significant enrichment of genes encoding enzymes involved in cholesterol, carbohydrate, and nucleotide metabolism. Within cluster 12, cluster 64 (124 genes, 1%) was enriched in genes with the Srebf binding motif. Srebf1c is stimulated by insulin, and this transcription factor regulates the gene expression of cholesterol metabolic enzymes (*28*). Indeed, this cluster contained 18 out of 37 cholesterol metabolism genes, and the genes transiently increased only in WT mice. Cluster 64 also included *Acly* and fatty acid desaturase 1 (*Fads1*), genes encoding the enzymes related to fatty acid synthesis.

Cluster 4, which had higher average expression in *ob/ob* mice than in WT mice, exhibited significant enrichment of lipid and carbohydrate metabolism, consistent with the high concentration of carbohydrate and triglyceride in *ob*/*ob* mice (Fig. 2 and fig. S4). This cluster also showed significant enrichment of the p53 binding motif. Cluster 4 included cluster 13 (988 genes, 7%), which had genes with higher expression in *ob*/*ob* mice and increased expression in the *ob/ob* mice after glucose administration. This cluster also showed enrichment in genes encoding enzymes involved in carbohydrate metabolism: genes encoding enzymes of glycolysis and the related pathway, such as phosphofructokinase, liver (*Pfkl*), citrate synthase (*Cs*), and pyruvate carboxylase (*Pcx*). This cluster also included genes encoding enzymes in the fatty acid synthesis pathway, such as acetyl-CoA carboxylase alpha (*Acaca*), *Acacb*, and *Fasn*, and the triglyceride synthesis pathway such as *Gpd1*, glycerol-3-phosphate acyltransferase, mitochondrial (*Gpam*), and glycerol-3-phosphate acyltransferase 4 (*Gpat4*). Although *Fasn* increased in WT mice and *ob*/*ob* mice, *Acacb*, *Cs*, *Gpat4*, and *Gpd1* increased only in *ob*/*ob* mice, indicating *ob*/*ob*-specific activation of lipid synthesis. This cluster was significantly enriched with the HBP1 binding motif and the HIC1 binding motif.

The genes in cluster 6 (1488 metabolites, 10%) showed no differences in expression between WT mice and *ob*/*ob* mice. However, they increased after glucose administration in both WT and *ob*/*ob* mice. This cluster exhibited significant enrichment of genes encoding proteins involved in protein processing in the endoplasmic reticulum and the spliceosome, and the genes were enriched for the XBP1 binding motif and the HIF1A binding motif.

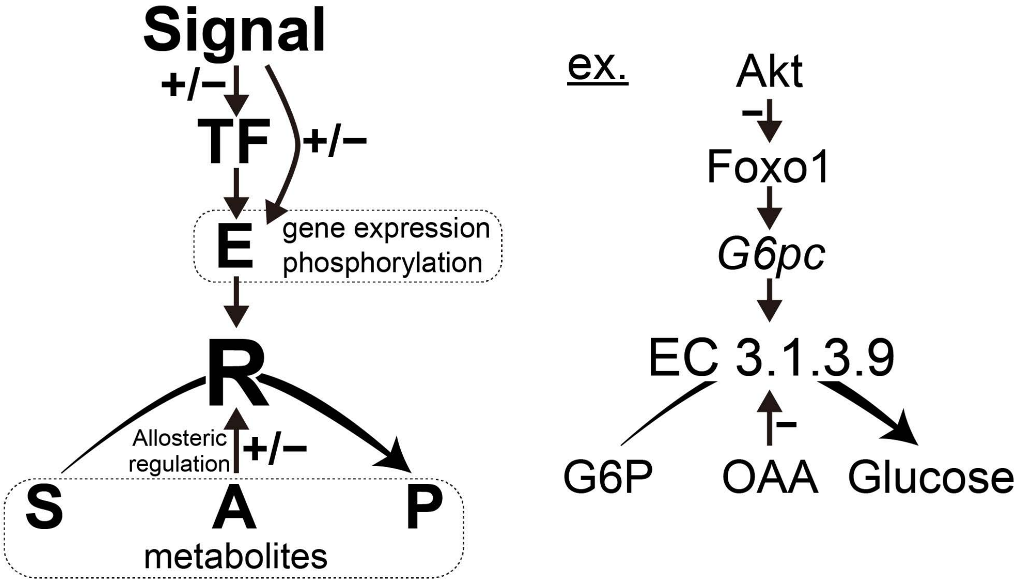


**Figure S1. Regulatory network for metabolic reactions.** (Left) A generic metabolic reaction (R) is catalyzed by metabolic enzyme (E) and involves metabolites that function as the substrate (S), product (P), or allosteric regulator (A). For reversible reactions, the product is also a substrate and the substrate is also a product (not shown). Positive and negative signs indicate positive and negative regulation, respectively. Regulation of a metabolic reaction by a metabolic enzyme consists of regulation by changing the amount of enzyme through gene expression and regulation by changing enzyme activity through posttranslational modifications, in particular phosphorylation. Gene expression is regulated by one or more transcription factors (TFs) and signaling molecules (Signals) regulate both transcription factor activity and metabolic enzyme activity by changing phosphorylation status. (Right) The metabolic reaction EC 3.1.3.9 is regulated through changes in expression of the gene encoding glucose-6-phosphatase (G6pc) and changes in the abundance of the substrate glucose-6-phosphate (G6P) and the allosteric inhibitor oxaloacetate (OAA). Expression of the *G6pc* gene is stimulated by Forkhead box protein O1 (FoxO1), and the activity of FoxO1 is negatively regulated by Akt through phosphorylation.

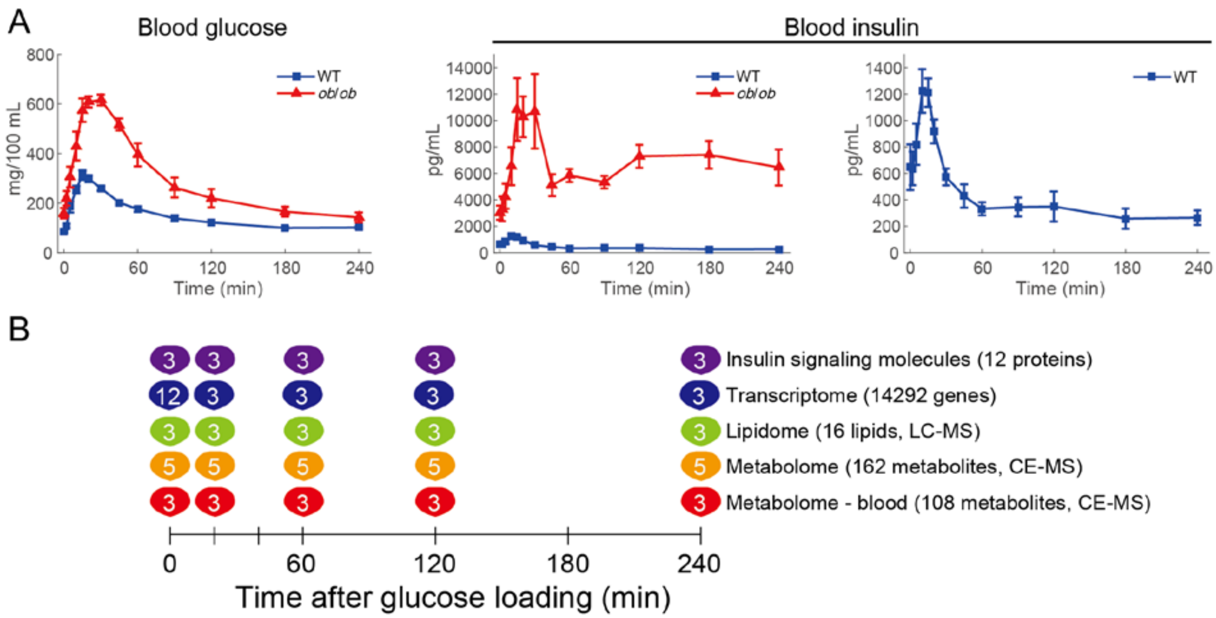


**Figure S2. Oral glucose administration and multi-omic measurements.** (**A**) Blood glucose and blood insulin of WT mice (blue) and *ob*/*ob* mice (red) during oral glucose administration. The means and SEMs of 5 mice are shown. (**B**) We orally administered glucose to 16 h-fasting WT and *ob*/*ob* mice, and collected the liver and blood at 0, 20, 60, 120, 240 min after administration. We performed metabolomics, transcriptomics, and western blotting for the phosphorylation of insulin signal molecules in the liver, and metabolomics in the blood. The number of the mice in each measurement is shown at each time point.

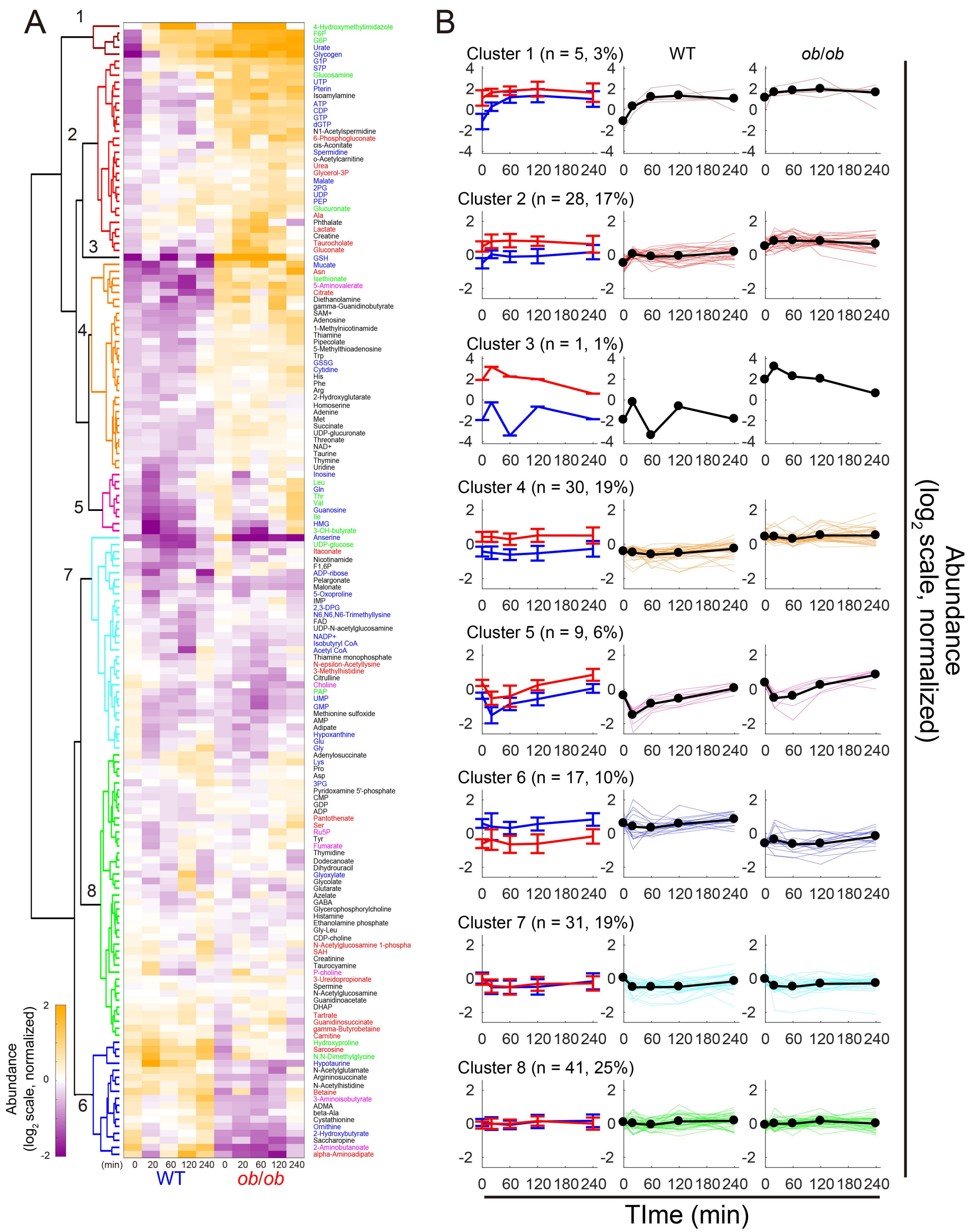


**Figure S3. Hierarchical clustering of time courses of metabolites in the liver.** (**A**) The heat map and hierarchical clustering of the time courses of metabolites in the livers of WT and *ob*/*ob* mice following oral glucose administration. The colors of and numbers on tree diagram indicate the cluster of each metabolite. To investigate the changes from fasting state, two time courses of each metabolite were divided by the geometric mean of the values of WT mice and *ob*/*ob* mice in fasting state (0 min), and then $\log_2$-transformed. The colors of the names of metabolites indicate WT mice-specific glucose-responsive metabolites (blue), *ob*/*ob* mice-specific glucose-responsive metabolites (red), common glucose-responsive metabolites (green), glucose-responsive metabolites showing opposite responses between WT mice and *ob*/*ob* mice (pink), and metabolites that are not glucose-responsive (black). (**B**) Averaged time courses of the metabolites for all 8 clusters. Left panel shows averaged time courses of the metabolites as the mean and standard deviation in a cluster for WT mice (blue) and *ob*/*ob* mice (red). Middle panel (WT mice) and right panel (*ob*/*ob* mice) show average (thick line) and individual (thin line) time courses of the metabolites in a cluster in WT or *ob/ob* mice.

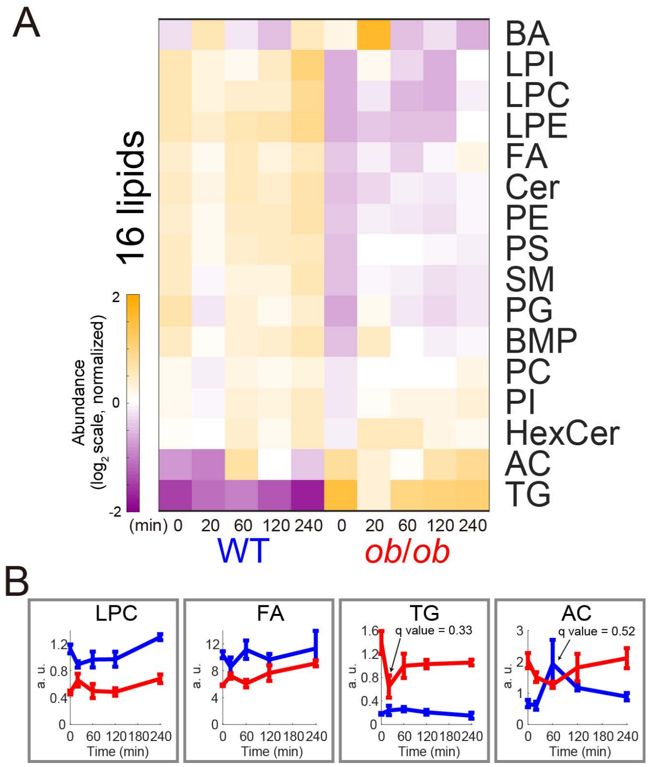


**Figure S4. Time courses of lipidomic data in the liver.** (**A**) The heat map of the time courses of 16 lipids in the livers of WT and *ob*/*ob* mice during oral glucose administration. Two time courses of each metabolite were divided by the geometric mean of the values of WT mice and *ob*/*ob* mice in fasting state (0 min), and then $\log_2$-transformed. Lipids were ordered by hierarchical clustering using Euclidean distance and Ward's method. No lipids showed significant change from fasting state (0 min) by oral glucose administration. The unabbreviated name of lipids can be found in table S2. (**B**) Time courses of the indicated lipids in the liver of WT mice (blue) and *ob*/*ob* mice (red) during oral glucose administration. Data are shown as the mean and SEM of 3 mice.

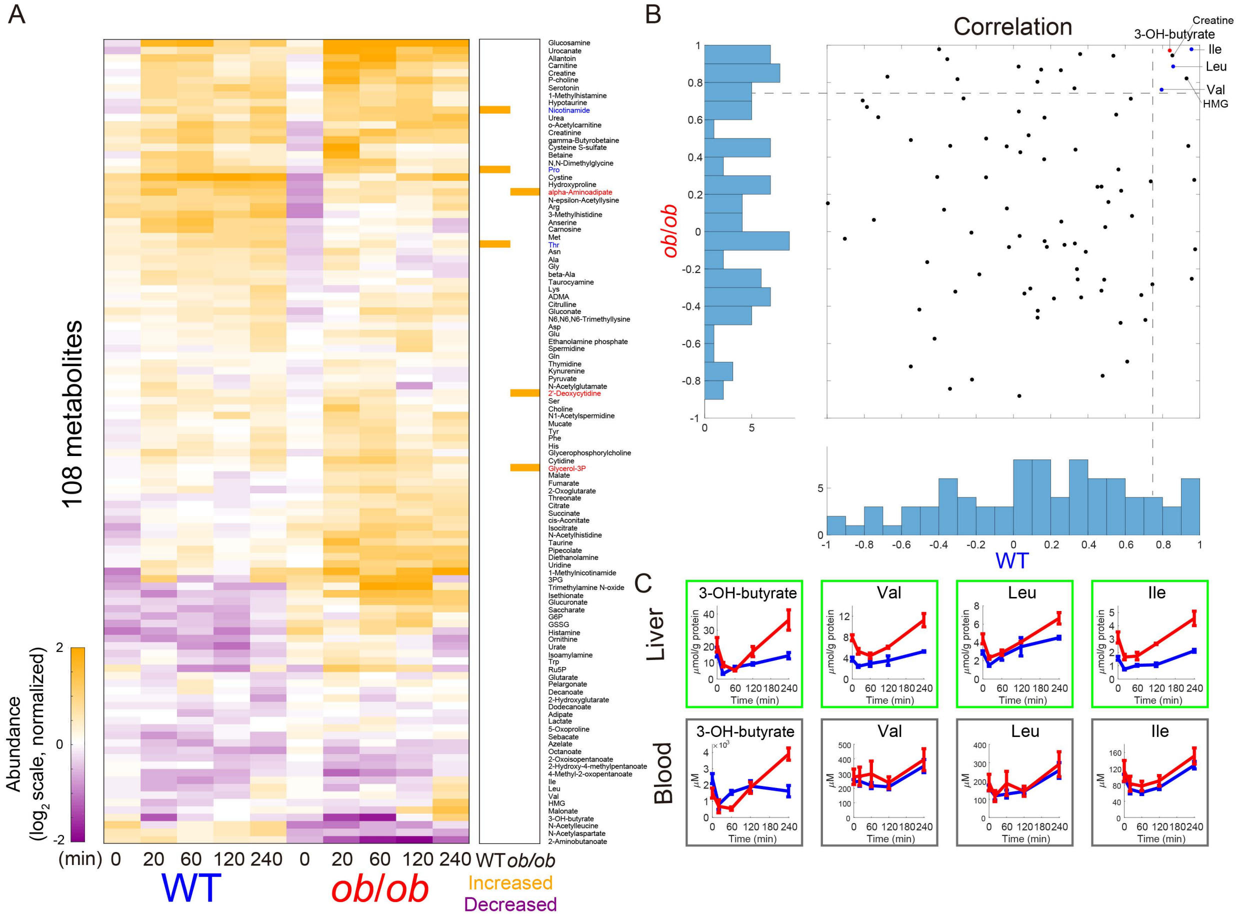


**Figure S5. Time courses of metabolites in the blood.** (**A**) Left: The heat map of the time courses of 108 metabolites in the blood of WT and *ob*/*ob* mice following oral glucose administration. Two time courses of each metabolite were divided by the geometric mean of the values of WT mice and *ob*/*ob* mice in fasting state (0 min), and then $\log_2$-transformed. Metabolites were ordered by hierarchical clustering using Euclidean distance and Ward's method. Right: The increased metabolites (orange) and unchanged metabolites (white) in WT mice and *ob*/*ob* mice. The changes from fasting state were determined by fold change and statistical test. The colors of the names of metabolites indicate WT mice-specific glucose-responsive metabolites (blue), *ob*/*ob* mice-specific glucose-responsive metabolites (red), and metabolites that were no responsive to glucose (black). (**B**) Histograms and scatter plot of Pearson's correlation coefficients between the time courses of metabolites measured in liver and blood in WT mice and *ob*/*ob* mice. Dashed lines correspond to correlation coefficients equal to 0.75. (**C**) Time courses of the indicated metabolites in the liver and blood of WT mice (blue) and *ob*/*ob* mice (red) following oral glucose administration. The indicated metabolites showed high correlations (correlation coefficient > 0.75) between the liver and the blood in both WT and *ob*/*ob* mice. The means and SEMs of mice are shown (n = 5 for liver, n = 3 for blood). The colors of the frames indicate common glucose-responsive metabolites (green) and non-responsive metabolites (gray).

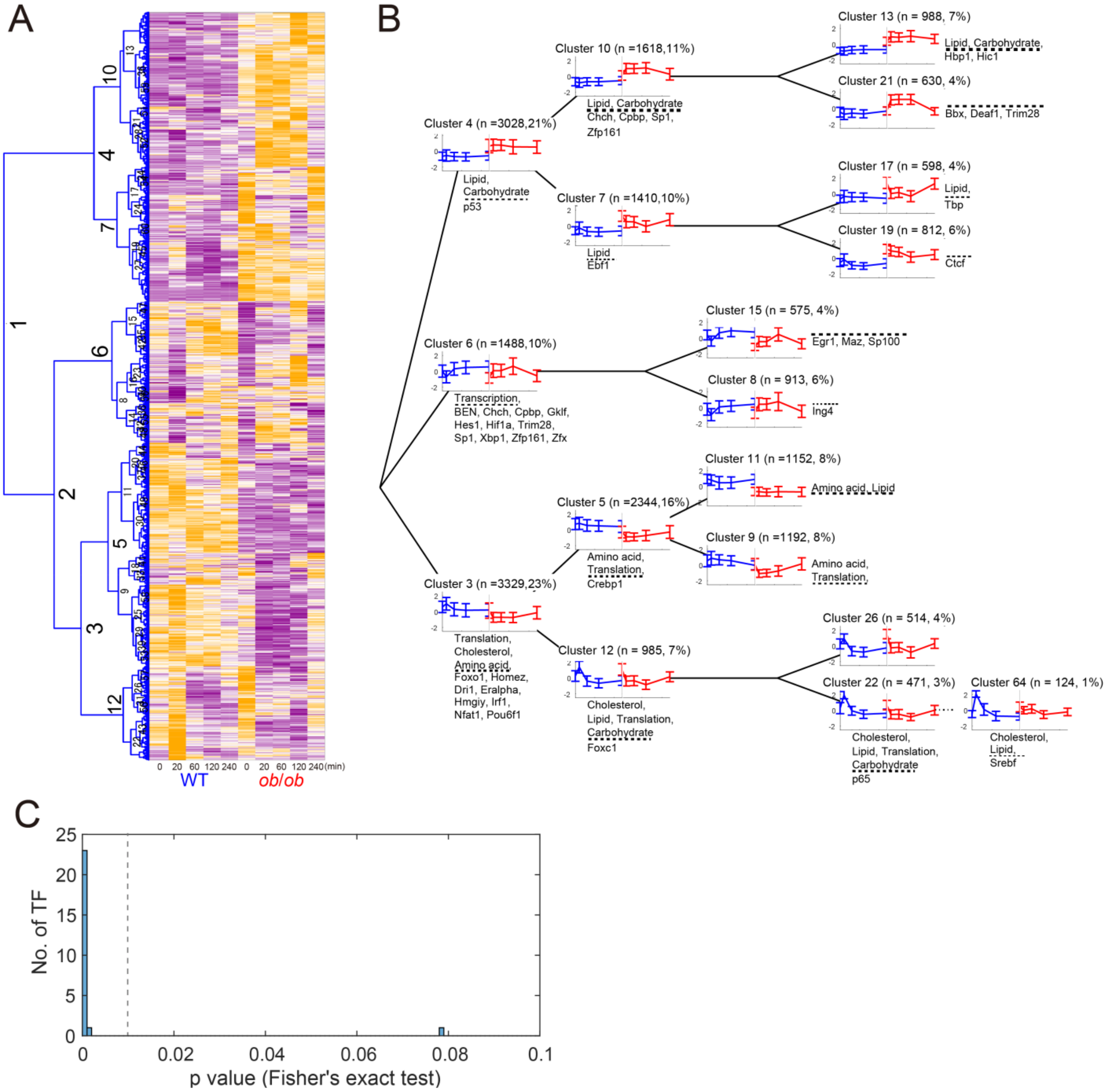


**Figure S6. Hierarchical clustering of time courses of gene expression in the liver and inference of regulatory connections between TFs and genes.** (**A**) The heat map and hierarchical clustering of the Z-score normalized time courses of gene expressions in the liver of WT and *ob*/*ob* mice following oral glucose administration. The hierarchical clustering was performed using Euclidean distance and Ward's method. The numbers on the tree diagram indicates the cluster identity. Each cluster includes only the genes that show a significant response at any time point either in WT mice or *ob*/*ob* mice or significant differences between WT mice and *ob*/*ob* mice at any time point (see Materials and Methods: Clustering analysis). These genes were determined by statistical test. (**B**) The averaged time courses of the gene expressions for each cluster of WT mice (blue) and *ob*/*ob* mice (red). The mean and standard deviation of the time courses of gene expressions in the cluster are shown. The time courses are presented on the tree diagram of hierarchical clustering. Significantly enriched pathways (above dashed line, p value < 0.01) and transcription factor motifs (below dashed line, q value < 0.1) in the cluster are described with the time courses. According to the enriched transcription factor motifs, we defined the regulatory connections between the transcription factors and the genes in the cluster. To avoid overestimation, we examined the enrichment of transcription factor binding motif in two children clusters of a cluster, and excluded the parent cluster from the inference if the transcription factor binding motif was significantly enriched in one of the two children clusters. The remaining transcription factor motifs, but not the excluded transcription factor motifs, are described here (above dashed line). The transcription factor motifs enriched in the upstream clusters are not described in the downstream clusters. Metabolic pathways described here are carbohydrate metabolism, amino acid metabolism, lipid metabolism, and cholesterol metabolism (below dashed line). Cholesterol metabolism is indicated by terpenoid backbone biosynthesis and steroid biosynthesis. (**C**) The histogram of the p values for the overlaps between the inferred genes of the transcription factors and those predicted from ChIP data. The ChIP data were obtained from the ChIP-Atlas database (*23*). The p values were calculated by one-tailed Fisher's exact test.

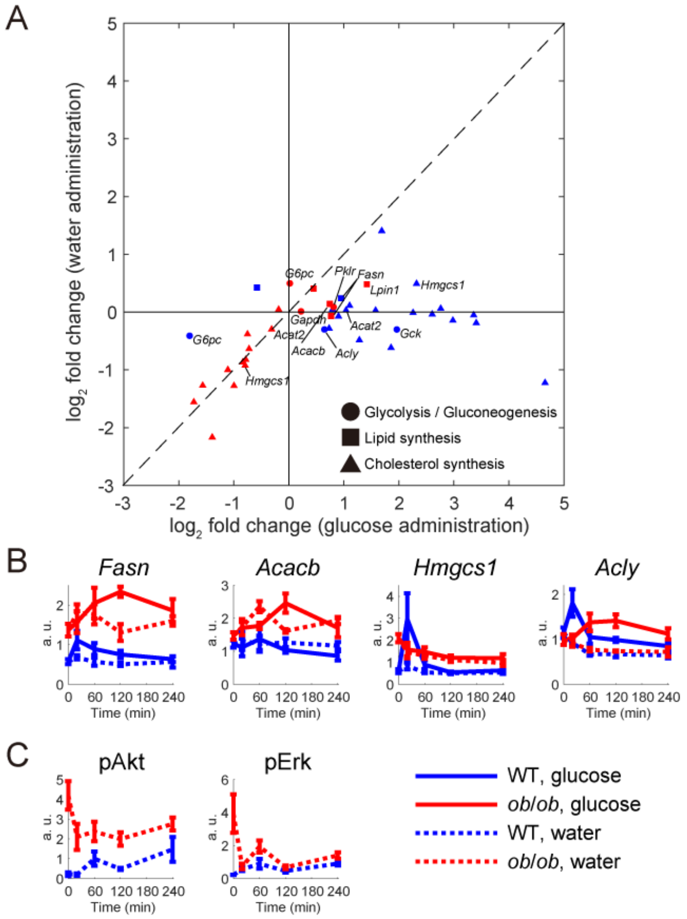


**Figure S7. Responses of glucose-responsive molecules during oral water administration.** (**A**) Responses of glucose-responsive genes to oral glucose administration and oral water administration in WT mice (blue) and *ob*/*ob* mice (red) measured by RT-PCR. The fold change of the mean amount at a significantly changed time point over the mean amount in fasting state was plotted for each gene. Genes near dashed line indicate that their expression following water administration was similar to the expression of those to glucose administration. The shapes of dots indicate the glucose-responsive genes in glycolysis and gluconeogenesis (circle), lipid synthesis (square), and cholesterol synthesis (triangle) (table S7). The significantly changed time points were determined from transcriptomic data. If a glucose-responsive gene had multiple significantly changed time points, only the maximum fold change was plotted for a upregulated gene, and only the minimum fold change was plotted for a downregulated gene. (**B**) Time courses of the indicated changes in gene expression in the liver of WT mice and *ob*/*ob* mice following oral glucose administration and water glucose administration measured by RT-PCR. The means and SEMs of mice are shown (n = 5 for glucose administration, n = 3 for water administration). (**C**) Time courses of the phosphorylation of Akt and Erk in the liver of WT mice and *ob*/*ob* mice following oral water administration. The means and SEMs of 3 mice are shown.

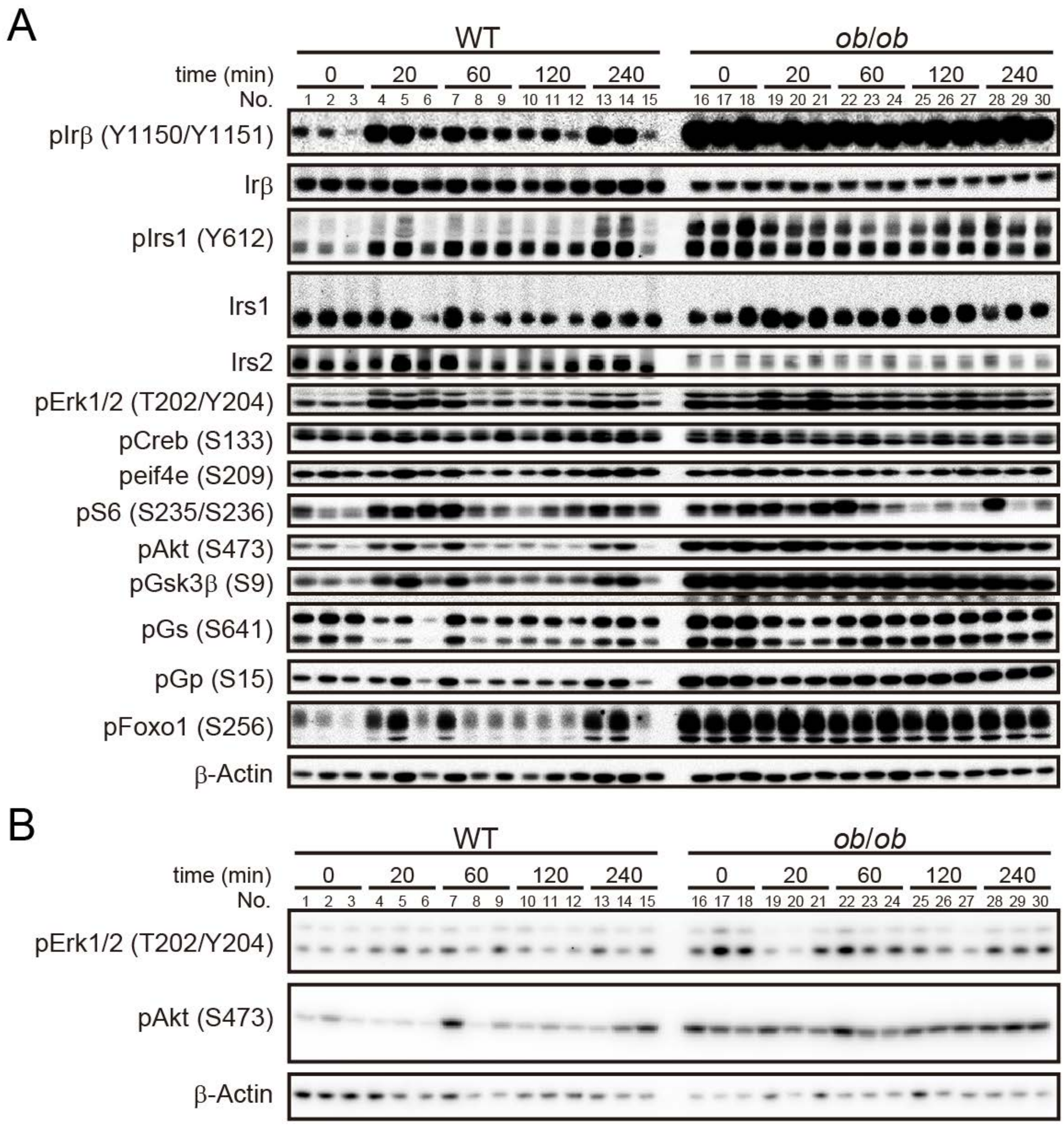


**Figure S8. Western blotting for insulin signaling molecules.** The amount and phosphorylation of the indicated insulin signaling molecules in the liver of WT mice and *ob*/*ob* mice at the indicated time point after oral glucose administration (**A**) and oral water administration (**B**). Residues in parentheses indicate the phosphorylation site(s) (human sequence numbering) recognized by the antibodies. All Western blot data for 3 mice are shown. In panel A, Lanes 5, 7, 13 and 14 were excluded in the following analysis, because the β-actin loading control for each indicated inconsistency from the other WT samples. The unabbreviated name of the insulin signaling molecules can be found in table S10.

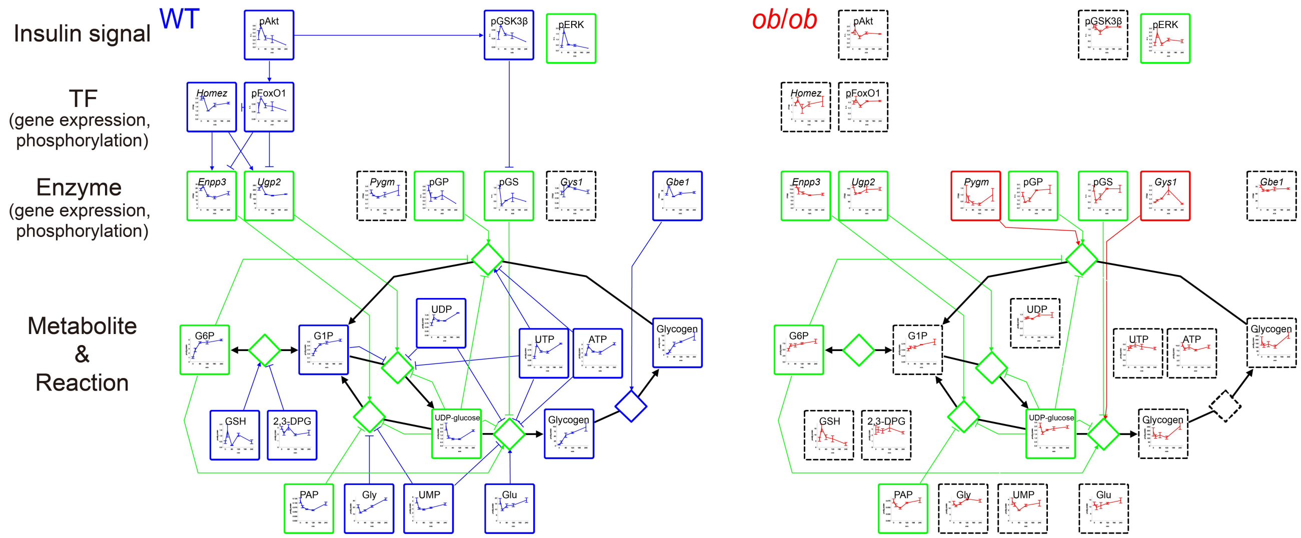


**Figure S9. The regulatory trans-omic network for glucose-responsive metabolic reactions in glycogen metabolism.** The regulatory trans-omic network for glucose-responsive metabolic reactions in glycogen metabolism in the liver of WT mice and *ob*/*ob* mice. The information for glycogen metabolism was obtained from "starch and sucrose metabolism" (mmu00500) in the KEGG database (*24*, *25*). Graphs of the time courses of measured molecules are shown for corresponding nodes as the means and SEMs of mice (n = 5 for metabolite, n = 11 or 12 for gene expression at 0 min, n = 3 for gene expression at 20 min, 60 min, 120 min, 240 min, n = 3 for phosphorylation). The colors of the frames indicate WT mice-specific glucose-responsive molecules (blue), *ob*/*ob* mice-specific glucose-responsive molecules (red), and common glucose-responsive molecules (green). The dashed frames indicate molecules that were not included in the glucose-responsive trans-omic network. Diamond nodes indicate metabolic reactions. The colored edges indicate inter-layer regulatory connections: WT mice-specific regulatory connections (blue), *ob*/*ob* mice-specific regulatory connections (red), and common inter-layer regulatory connections (green). From metabolite to reaction, only allosteric regulatory connections are colored. Black edges indicate the relationship between metabolic reactions and its substrate/product. The reversibility of metabolic reactions was obtained from the KEGG database (*24*, *25*).

In glycogen metabolism, G1P and glycogen increased specifically in WT mice, and UDP-glucose decreased in both WT and *ob*/*ob* mice during glucose administration. The decrease of pGs activated the consumption of UDP-glucose and the synthesis of glycogen in both WT and *ob*/*ob* mice. UDP-glucose pyrophosphorylase 2 (*Ugp2*), the downregulated gene in both mice, was also identified as the glucose-responsive molecules related to the decrease of UDP-glucose. In WT mice, glucose-responsive molecules contributing to the increase of glycogen were pGs, pGp, glucan (1,4-alpha-), branching enzyme 1 (*Gbe1*), G6P, ATP, UDP-glucose, and UMP. Among these glucose-responsive molecules, the WT mice-specific responses were the upregulation of *Gbe1*, the increase of ATP, the decrease of UMP. G6P was a common glucose-responsive metabolite, but the amplitude was quite larger in WT mice than in *ob*/*ob* mice (fold change at 60 min after glucose administration = 4.6, 1.4 in WT and *ob*/*ob* mice, respectively), indicating that the responses of G6P also might contribute to WT mice-specific increase of glycogen.

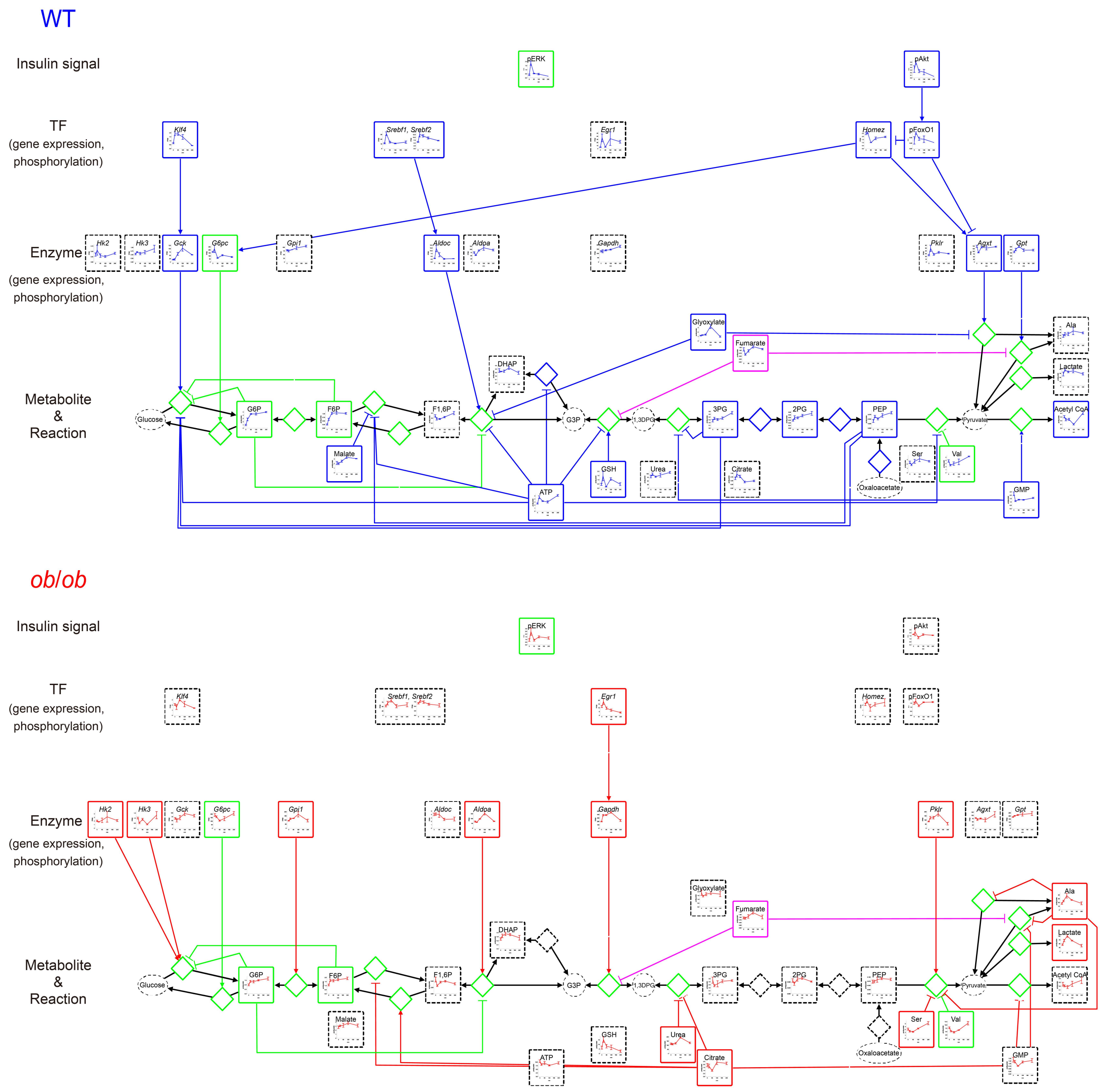


**Figure S10. The regulatory trans-omic network for glucose-responsive metabolic reactions in glycolysis and gluconeogenesis.** The regulatory trans-omic network for glucose-responsive metabolic reactions in glycolysis and gluconeogenesis in the liver of WT mice and *ob*/*ob* mice. The information of glycolysis and gluconeogenesis were obtained from “glycolysis / gluconeogenesis” (mmu00010) in the KEGG database (*24*, *25*). The glucose-responsive gene *Pgk1-rs7* is excluded. Graphs of the time courses of measured molecules are shown for corresponding nodes as the means and SEMs of mice (n = 5 for metabolite, n = 11 or 12 for gene expression at 0 min, n = 3 for gene expression at 20 min, 60 min, 120 min, 240 min, n = 3 for phosphorylation). The colors of the frames indicate WT mice-specific glucose-responsive molecules (blue), *ob/ob* mice-specific glucose-responsive molecules (red), common glucose-responsive molecules (green), and molecules showing opposite responses between WT mice and *ob/ob* mice (pink). The dashed frames indicate molecules that were not included in the glucose-responsive trans-omic network. Diamond nodes indicate metabolic reactions. The colored edges indicate inter-layer regulatory connections: WT mice-specific regulatory connections (blue), *ob*/*ob* mice-specific regulatory connections (red), common inter-layer regulatory connections (green), and opposite inter-layer regulatory connections (pink). From metabolite to reaction, only allosteric regulatory connections are colored. Black edges indicate the relationship between metabolic reactions and its substrate/product. The reversibility of metabolic reactions was obtained from the KEGG database (*24*, *25*).

In glycolysis and gluconeogenesis, G6P and F6P increased in both WT and *ob*/*ob* mice, 3-phosphoglyceric acid (3PG), 2PG, and PEP increased specifically in WT mice, Acetyl CoA decreased specifically in *ob*/*ob* mice, and Ala and lactate increased specifically in *ob*/*ob* mice. In WT mice, the gene encoding glucokinase (*Gck*) was strongly upregulated (fold change at 60 min = 3.5), and *G6pc* was strongly downregulated (fold change at 60 min = 0.39), both of which can account for the large increase of G6P (fold change at 60 min = 4.6). In contrast, both changes of gene expression were weaker in *ob*/*ob* mice (fold change at 60 min = 1.1, 0.57, respectively), consistent with the small increase of G6P (fold change at 60 min = 1.4).

ATP, 3PG, PEP, malate, and glyoxylate increased during oral glucose administration, and inhibited glucose-responsive metabolic reactions in glycolysis/gluconeogenesis specifically in WT mice. Especially, ATP was identified as the inhibitory allosteric regulator of six glucose-responsive metabolic reactions in glycolysis/gluconeogenesis in the WT mice network. For example, the conversion of F6P to F1,6P was allosterically inhibited by ATP. We also confirmed that, according to the BRENDA database (*26*), this metabolic reaction was not saturated by ATP (saturation index = 0.12) (fig. S13 and table S13).

In *ob*/*ob* mice, some of glycolytic enzyme genes such as glucose phosphate isomerase 1 (*Gpi*), aldolase A (*Aldoa*), *Gapdh*, and *Pklr* were upregulated specifically in *ob*/*ob* mice, most of which showed slow responses. p53 was identified as the regulator of these *ob*/*ob* mice-specific glucose-responsive genes. In addition, inhibitory allosteric regulations were absent in *ob*/*ob* mice. Rather, because citrate decreased specifically in *ob*/*ob* mice, three inhibitory regulations of metabolic reactions in glycolysis/gluconeogenesis by citrate were weakened during oral glucose administration. These regulations may account for the *ob*/*ob* mice-specific increase of lactate.

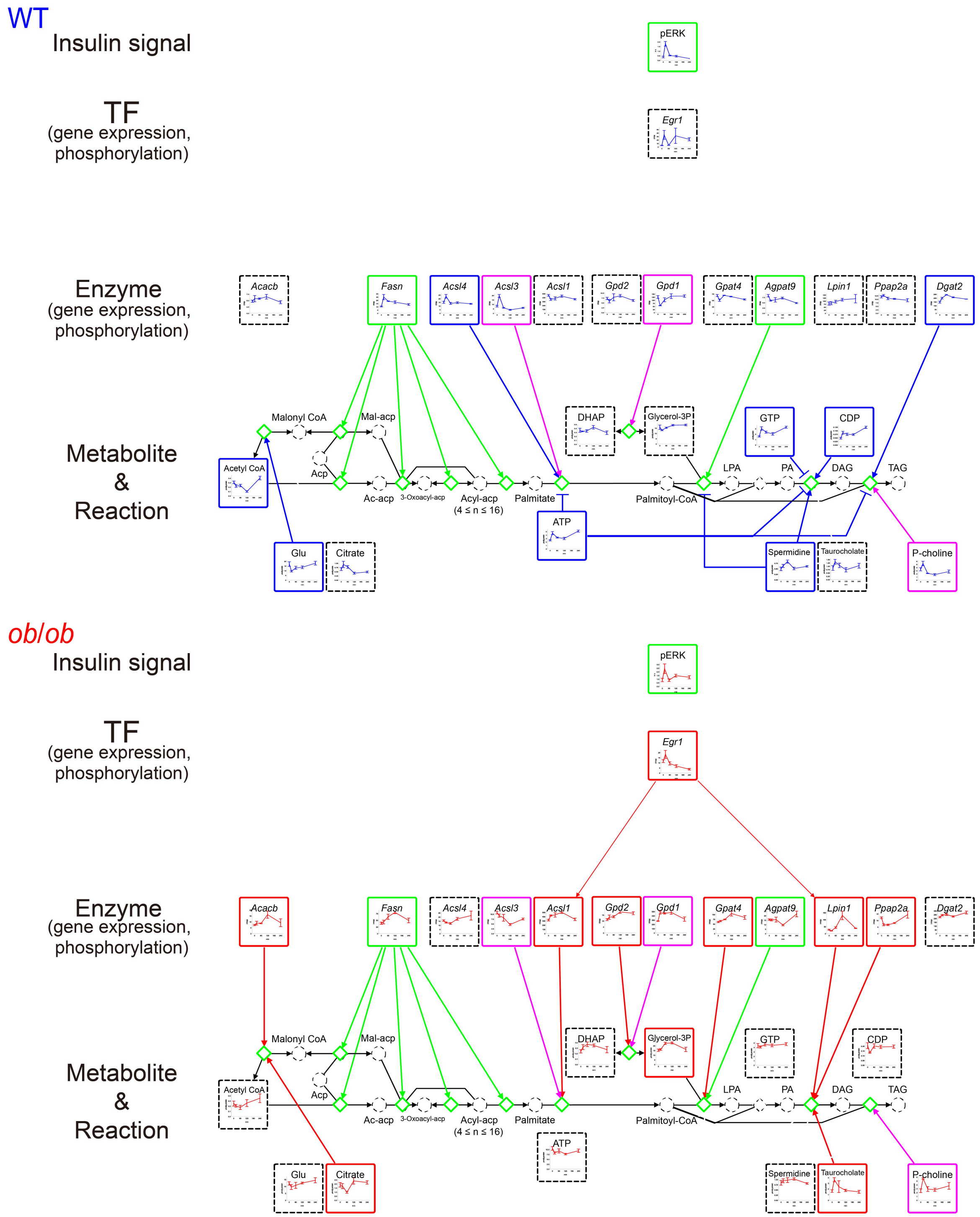


**Figure S11. The regulatory trans-omic network for glucose-responsive metabolic reactions in lipid synthesis.** The regulatory trans-omic network for glucose-responsive metabolic reactions in lipid synthesis in the liver of WT mice and *ob*/*ob* mice. The information of lipid synthesis was obtained from “fatty acid biosynthesis” (mmu00061), “glycerolipid metabolism” (mmu00561), and “glycerophospholipid metabolism” (mmu00564) in the KEGG database (*24*, *25*). Some intermediates in fatty acid biosynthesis are not plotted. Graphs of the time courses of measured molecules are shown for corresponding nodes as the means and SEMs of mice (n = 5 for metabolite, n = 11 or 12 for gene expression at 0 min, n = 3 for gene expression at 20 min, 60 min, 120 min, 240 min, n = 3 for phosphorylation). The colors of the frames indicate WT mice-specific glucose-responsive molecules (blue), *ob/ob* mice-specific glucose-responsive molecules (red), common glucose-responsive molecules (green), and molecules showing opposite responses between WT mice and *ob/ob* mice (pink). The dashed frames indicate molecules that were not included in the glucose-responsive trans-omic network. Diamond nodes indicate metabolic reactions. The colored edges indicate inter-layer regulatory connections: WT mice-specific regulatory connections (blue), *ob*/*ob* mice-specific regulatory connections (red), common inter-layer regulatory connections (green), and opposite inter-layer regulatory connections (pink). From metabolite to reaction, only allosteric regulatory connections are colored. Black edges indicate the relationship between metabolic reactions and its substrate/product. The reversibility of metabolic reactions was obtained from the KEGG database (*24*, *25*).

In the genes in lipid synthesis, *Fasn* was upregulated in both WT and *ob*/*ob* mice, three genes were upregulated specifically in WT mice, such as long-chain acyl-CoA synthetases (*Acsl3* and *Acsl4*), and six genes were upregulated specifically in *ob*/*ob* mice, such as *Acacb*, *Gpd1*, *Gpat4*, and lipin 1 (*Lpin1*). p53 and *Egr1* were identified as the transcription factor regulating these *ob*/*ob* mice-specific glucose-responsive genes. In WT mice, 5 out of 7 allosteric regulations by glucose-responsive metabolites, including increased ATP, inhibited the metabolic reactions in lipid synthesis. In contrast, increased taurocholate and phosphorylcholine (P-choline) allosterically activated metabolic reactions in lipid synthesis. These results suggest that lipid synthesis is activated through gene expression and allosteric mechanisms in *ob*/*ob* mice.

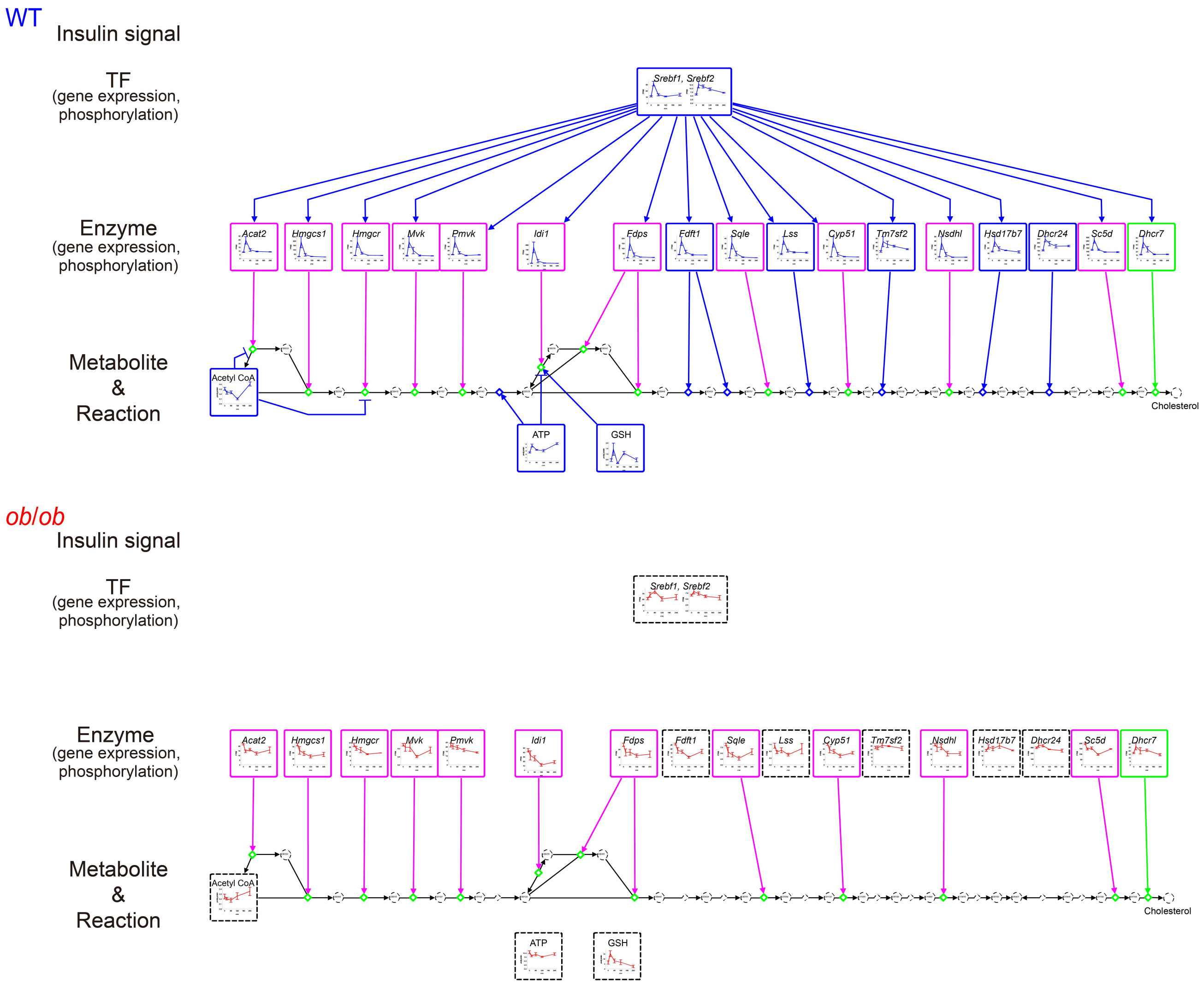


**Figure S12. The regulatory trans-omic network for glucose-responsive metabolic reactions in cholesterol synthesis.** The regulatory trans-omic network for glucose-responsive metabolic reactions in cholesterol synthesis in the liver of WT mice and *ob*/*ob* mice. The information of cholesterol synthesis were obtained from "terpenoid backbone biosynthesis" (mmu00900) and "steroid biosynthesis" (mmu00100) in the KEGG database (*24*, *25*). Graphs of the time courses of measured molecules are shown for corresponding nodes as the means and SEMs of mice (n = 5 for metabolite, n = 11 or 12 for gene expression at 0 min, n = 3 for gene expression at 20 min, 60 min, 120 min, 240 min, n = 3 for phosphorylation). The colors of the frames indicate WT mice-specific glucose-responsive molecules (blue), *ob/ob* mice-specific glucose-responsive molecules (red), common glucose-responsive molecules (green), and molecules showing opposite responses between WT mice and *ob/ob* mice (pink). The dashed frames indicate molecules that were not included in the glucose-responsive trans-omic network. Diamond nodes indicate metabolic reactions. The colored edges indicate inter-layer regulatory connections: WT mice-specific regulatory connections (blue), *ob*/*ob* mice-specific regulatory connections (red), common inter-layer regulatory connections (green), and opposite inter-layer regulatory connections (pink). From metabolite to reaction, only allosteric regulatory connections are colored. Black edges indicate the relationship between metabolic reactions and its substrate/product. The reversibility of metabolic reactions was obtained from the KEGG database (*24*, *25*).

In WT mice, most of the genes in cholesterol synthesis were upregulated transiently. *Srebf* was identified as the transcription factor regulating these glucose-responsive genes. Of the 17 genes, 11 genes were downregulated in *ob*/*ob* mice, all of which similarly decreased by oral water administration only in *ob*/*ob* mice (fig. S7). This result indicates that the responses of these genes are not specific to orally administrated glucose in *ob*/*ob* mice. WT mice-specific glucose-responsive metabolites also allosterically activated metabolic reactions in cholesterol synthesis, except for the regulation of EC 5.3.3.2 by ATP.

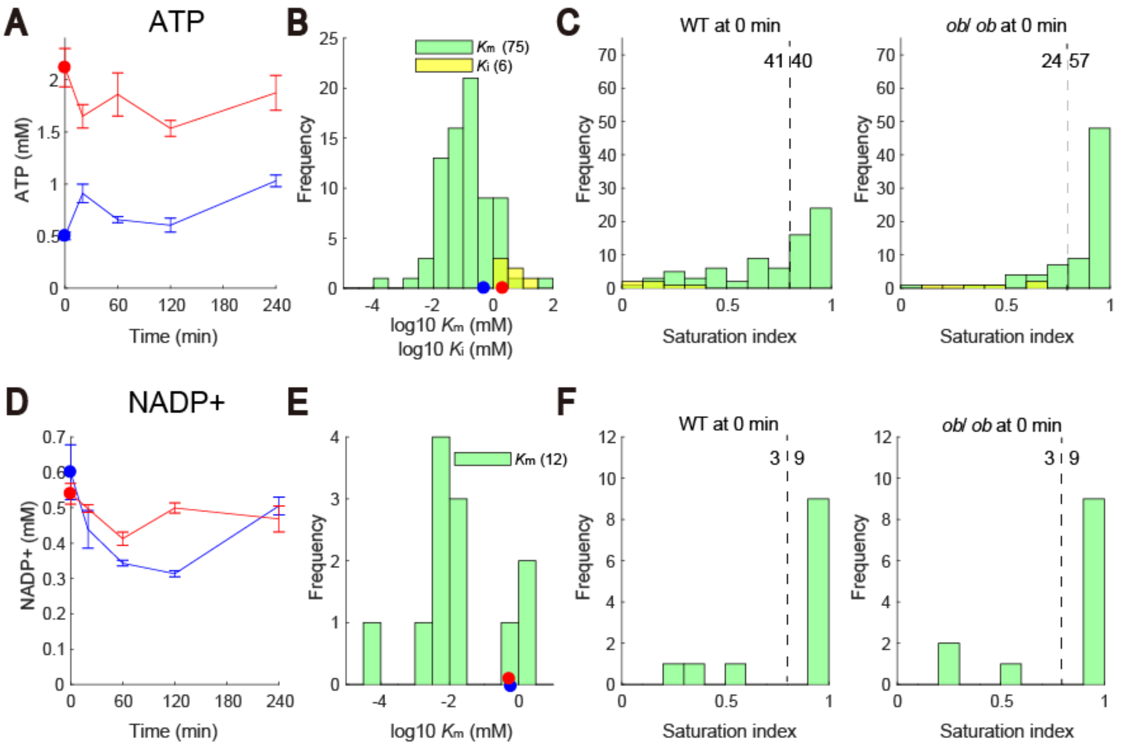


**Figure S13. $K_m$ and $K_i$ of metabolic reactions regulated by ATP and NADP+.** (**A**) Graph of the time courses for ATP in the liver of WT mice (blue) and *ob*/*ob* mice (red) following oral glucose administration. The means and SEMs of 5 mice are shown. The dots indicate the fasting values. (**B**) Histogram of $K_m$ values and $K_i$ values of metabolic reactions regulated by ATP according to the BRENDA database (*26*) (table S13). The dots indicate the fasting values of ATP in WT mice (blue) and *ob*/*ob* mice (red). (**C**) Histograms of the saturation indices of metabolic reactions regulated by ATP in WT mice and *ob*/*ob* mice in fasting state (0 min). The saturation indices of metabolic reactions regulated by ATP as substrate/product (green) were defined as [ATP] / [ATP] + $K_m$, and those as allosteric regulator (yellow) were defined as [ATP] / [ATP] + $K_i$. Saturation index closer to 1 means higher saturation, and that closer to 0 means lower saturation. We defined unsaturated metabolic reactions as metabolic reactions with saturation indices that were less than 0.8 (dashed line). The numbers besides the dashed lines indicate the numbers of unsaturated metabolic reactions (left) and saturated metabolic reactions (right). (**D**) Graph of the time courses for NADP+ in the liver of WT mice (blue) and *ob*/*ob* mice (red) following oral glucose administration. (**E**) Histogram of $K_m$ values of metabolic reactions regulated by NADP+ according to the BRENDA database (*26*) (table S13). $K_i$ values were not available in the BRENDA database. The dots indicate the fasting values of NADP+ in WT mice (blue) and *ob*/*ob* mice (red). (**F**) Histograms of the saturation indices of metabolic reactions regulated by NADP+ in WT mice and *ob*/*ob* mice in fasting state (0 min) with the numbers left of the dashed line indicating unsaturated metabolic reactions and right indicating saturated metabolic reactions.

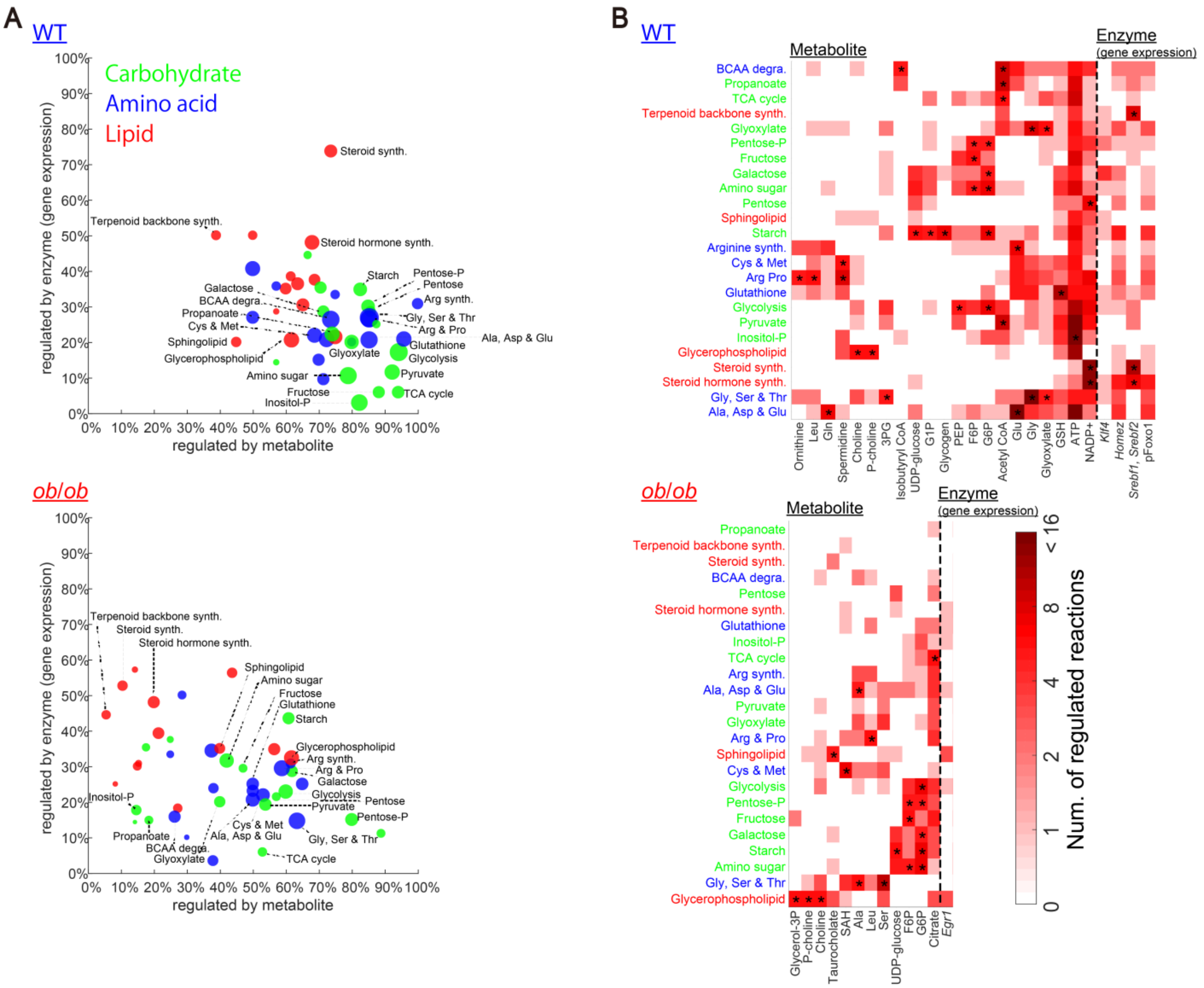

**Figure S14. Metabolic reactions regulated by glucose-responsive molecules in each metabolic pathway node.** (**A**) For each metabolic pathway node, the percent of regulated metabolic reactions by glucose-responsive metabolites (*x*-axis) and by glucose-responsive genes of metabolic enzyme (*y*-axis) is plotted for WT mice and *ob/ob* mice. The size of the dots indicates the number of regulated metabolic reactions in each metabolic pathway node either by glucose-responsive metabolites or genes or both. The colors of the dots indicate the classes of metabolic pathway node according to the KEGG database (*24*, *25*): carbohydrate (green), amino acid (blue), and lipid (red). (**B**) Heat maps showing the number of regulated metabolic reactions in each metabolic pathway node (rows) by each glucose-responsive metabolite (left columns) and each transcription factor-dependent glucose-responsive genes of metabolic enzymes (right columns) in WT mice and *ob/ob* mice. The signs * indicate the significant associations (p value < 0.05) between metabolic reactions in the metabolic pathway node and those regulated by the glucose-responsive molecules (table S14). The p values were calculated by one-tailed Fisher's exact test. Only metabolic pathway nodes with significant associations with any glucose-responsive molecule are shown. Only glucose-responsive metabolites with significant associations with any metabolic pathway node are shown.

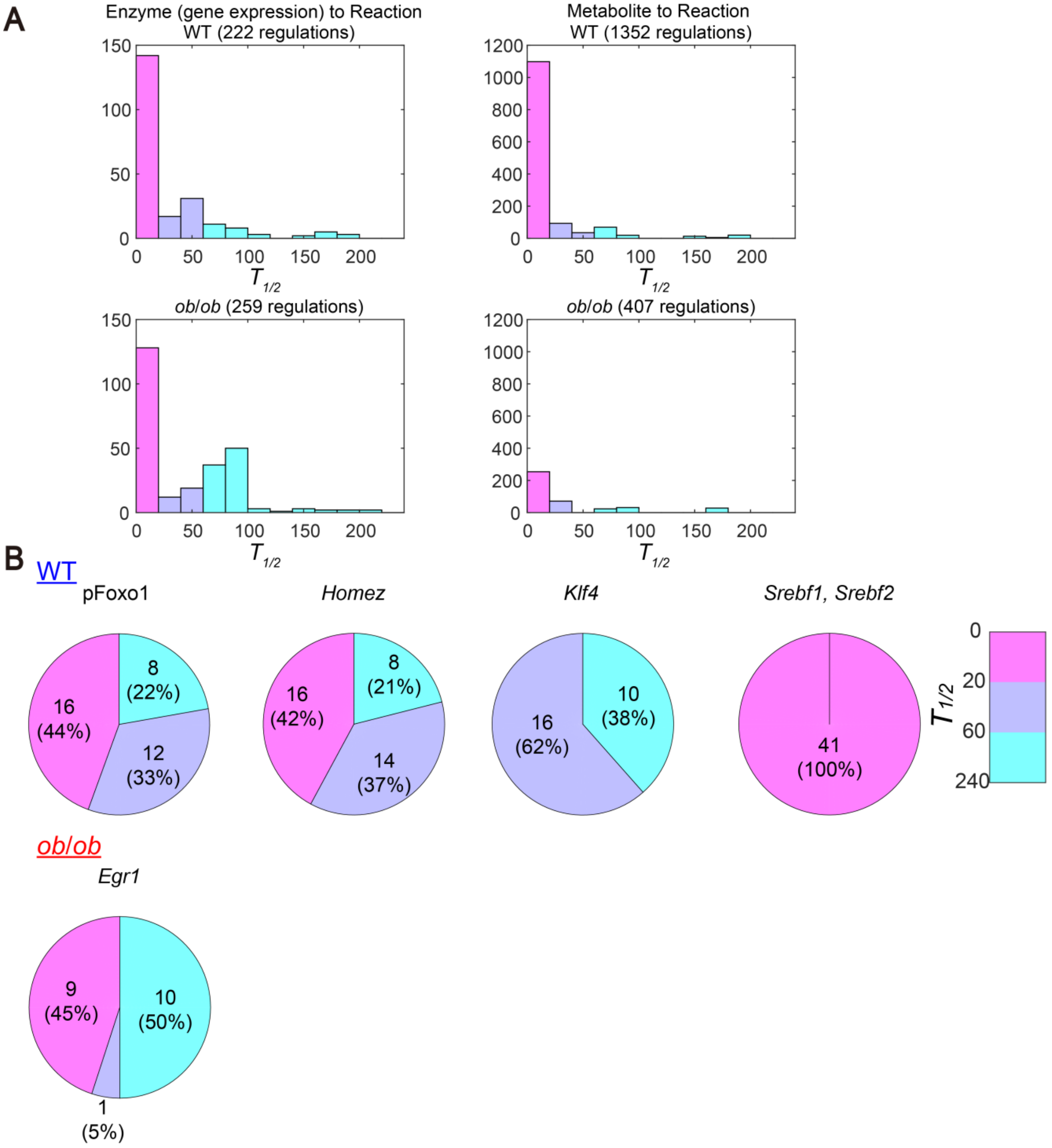


**Figure S15. $T_{1/2}$ values of the inter-layer regulatory connections from glucose-responsive molecules to metabolic reactions.** (**A**) The number of the inter-layer regulatory connections with indicated $T_{1/2}$ values from the Enzyme layer to the Reaction layer and from the Metabolite layer to the Reaction layer in WT mice and *ob*/*ob* mice. (**B**) The number and percentage of $T_{1/2}$ values of the inter-layer regulatory connections of metabolic reactions by the indicated transcription factor-dependent glucose-responsive genes encoding metabolic enzymes in WT mice and *ob/ob* mice. The colors of the pies represent the ranges of $T_{1/2}$ values as shown in the color bar on the right.

**Supplementary tables**

**Table S1: Metabolomic data in the liver**.
**Table S2: Lipidomic data**.
**Table S3: Metabolomic data in the blood**.
**Table S4: Transcriptomic data**.
**Table S5: Pathway enrichment analysis of glucose-responsive genes and genes showing the differences in the amounts of expression between WT mice and *ob*/*ob* mice before oral glucose administration**.
**Table S6: RT-PCR data.**
**Table S7: Enrichment analysis of gene clusters.**
**Table S8: Inferred regulatory connections between TFs and genes.**
**Table S9: Overlap between the inferred genes of TF and those predicted from experimental ChIP data.**
**Table S10: Western blotting data.**
**Table S11: Regulatory trans-omic network for glucose-responsive metabolic reactions.**
**Table S12: Enrichment analysis of downstream genes of each TF.**
**Table S13: Enzyme binding affinity $K_m$ and $K_i$.**
**Table S14: Significant associations between glucose-responsive molecules and metabolic pathways.**
